# Self-dyeing textiles grown from cellulose-producing bacteria with engineered tyrosinase expression

**DOI:** 10.1101/2023.02.28.530172

**Authors:** Kenneth T. Walker, Jennifer Keane, Vivianne J. Goosens, Wenzhe Song, Koon-Yang Lee, Tom Ellis

## Abstract

Environmental concerns are driving interests in post-petroleum synthetic textiles produced from microbial and fungal sources. Bacterial cellulose is a promising sustainable leather alternative, on account of its material properties, low infrastructure needs and biodegradability. However, for alternative textiles like bacterial cellulose to be fully sustainable, alternative ways to dye textiles need to be developed alongside alternative production methods. To address this, we here use genetic engineering of *Komagataeibacter rhaeticus* to create a bacterial strain that grows self-dyeing bacterial cellulose. Dark black pigmentation robust to material use is achieved through melanin biosynthesis in the bacteria from recombinant tyrosinase expression. Melanated bacterial cellulose production can be scaled up for the construction of prototype fashion products, and we illustrate the potential of combining engineered self-dyeing with tools from synthetic biology, via the optogenetic patterning of gene expression in cellulose-producing bacteria. With this work, we demonstrate that combining genetic engineering with current and future methods of textile biofabrication has the potential to create a new class of textiles.

## Main

The textile and leather industry impacts the environment – contributing to greenhouse gas emissions from agricultural production and industrial processing, water pollution through tanning and dyeing, and microplastic pollution from synthetic fibre shedding^1–3^. To reduce the impact of this industry, new sustainable biomaterials are under commercial development. These include mycelium and plant fibre based leather alternatives^4,5^. These endeavours are the successful result of combining biological production with engineering and chemical processing to refine these natural biomaterials into alternative textiles. However, the industry is yet to employ genetic engineering of these material producing organisms to take advantage of the sustainable methods biological systems use to enhance the physical and aesthetic properties of biomaterials.

The field of engineered living materials (ELMs) uses the tools of synthetic biology to reprogram living cells at the DNA level to build new or enhanced biomaterials for specific applications^6,7^. Bacterial cellulose (BC) is a promising natural biomaterial, produced most effectively by bacteria in the gram-negative genus *Komagataeibacter*^8^. In carbon rich media, these bacteria polymerise and secrete linear chains of glucose. These chains then self-assemble into a dense interconnected mesh of cellulose fibres. This cellulose mesh, called a pellicle, floats at the air:water interface and envelops and protects the growing cells, like a biofilm^9^. Key to the industrial interest in BC, it can be grown quickly, cheaply and sustainably — a BC pellicle can be grown in 7-14 days, in high yields (> 10 grams per litre) and from waste feedstocks, such as rotten fruit juice, glycerol and molasses^10–12^. Additionally, BC has advanced material properties such as high tensile strength, high water holding capacity and high purity^13,14^. These features have led to interest in using BC in high end acoustic devices, as a battery separator membrane and in wound healing^15–20^. The ease of growing BC has also led to BC becoming an attractive prototype biomaterial for some in design and fashion who seek to speculate on methods of sustainable textile production^21^. The production of BC by culturable, low-risk bacteria also makes BC accessible to those seeking to modify it genetically using synthetic biology. BC therefore represents an ideal ‘blank slate’ for ELM research.

BC ELM research has focused on the genetic engineering of *Komagataeibacter* and other organisms such as *Saccharomyces cerevisiae* that can be co-cultured with *Komagataeibacter*. The use of incorporated *S. cerevisiae* has allowed for the production of pellicles that can sense and respond to chemical and light stimuli^22^. To facilitate the genetic engineering of *Komagataeibacter* a modular gene cloning tool kit, the Komagataeibacter Toolkit (KTK), has been created and characterised using *Komagataeibacter rhaeticus*. Contributions to this synthetic biology tool kit include a selection of modular DNA parts, such as constitutive and inducible promoters, vectors and fluorescent markers^23–25^. Such parts have been used to make engineered *Komagataeibacter* that can produce alternative polymers, such as chitin, hyaluronic acid and curli fibres^25–27^. Additionally, engineered multicellular communication has been established in a pellicle, through cell-to-cell signalling between *K. rhaeticus* cells using quorum sensing molecules^28^. However, despite these achievements genetic engineering has yet to be used to further the development of BC as a sustainable biomaterial in textiles and fashion.

Biomaterials in nature, such as hair and skin, use incorporated cells to produce pigments that colour the biomaterial *in situ* in a low-impact sustainable manner. The parallel process used in industrial colouring of fabric materials, textile dyeing, requires chemical reactions and is highly damaging to the environment. Inspired by natural pigment production, we set out to engineer a self-dyeing BC material through the genetic engineering of *K. rhaeticus*. Black dye is one of the most consumed dyes in the world, and one of the most difficult to recreate using sustainable dyeing approaches^29,30^. We decided to engineer the biosynthesis of the dark melanin pigment, eumelanin, into *K. rhaeticus*. Eumelanin, a ubiquitous pigment found across biological kingdoms, is stable in high heat and over long timespans^31^. Crucially, eumelanin has low water solubility, a property shared by many common dyes, such as indigo, that contributes to the colour fastness of a pigment^32^. Additionally, eumelanin also offers several other interesting properties, such as electrical conductivity, broad-band light and UV absorption, and protection from ionising radiation^33–37^. We demonstrate here that the production of pigmented cellulose from *K. rhaeticus* can be produced at large enough quantities for the prototyping of fashion products. Furthermore, we illustrate the potential of combining melanin biosynthesis with other synthetic biology tools, through the optogenetic patterning of gene expression in growing BC pellicles.

## Results

### Melanin production from *K. rhaeticus*

Recombinant production of eumelanin has been demonstrated in *E. coli* and *Vibrio natrigens* in the pursuit of diverse applications like bioremediation, and bioelectronics^38–41^. The bacterial production of eumelanin requires only a single enzyme – tyrosinase – that catalyses the oxidation of L-tyrosine to dopaquinone — the rate limiting step in eumelanin synthesis. In oxygenated and temperate conditions, dopaquinone spontaneously converts to eumelanin via several steps **(Fig. 1a**). The prokaryotic tyrosinases that have been tested in a recombinant context are, MelA from *Rhizobium etli* and Tyr1 from *Bacillus megaterium*^40,42^. We decided to focus on Tyr1 for this study, due to its smaller size and proven use in non-model organisms. Using our KTK system for modular cloning, we created two constitutive *K. rhaeticus* Tyr1 expression strains: the plasmid-based *K. rhaeticus* _*p*_*tyr1* and the chromosomally-integrated *K. rhaeticus tyr1* (**Fig. 1b**). Both strains used identical upstream and downstream DNA parts around the *tyr1* coding sequence, which included the synthetic constitutive promoter pJ23104, which was previously found to have the strongest expression strength of a library of promoters characterised in *K. rhaeticus*^25^.

**Figure 1.**
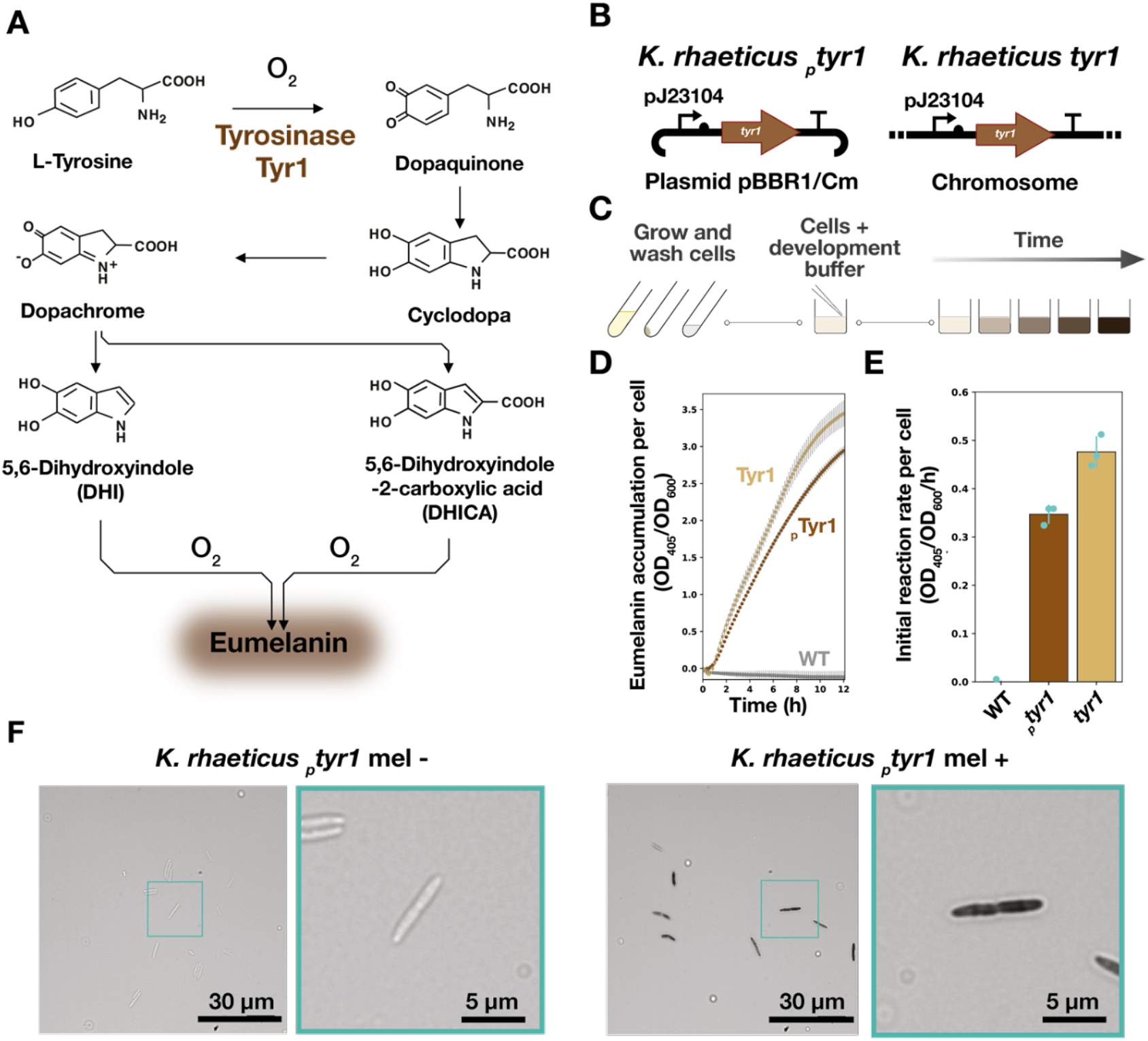
Eumelanin production from *K. rhaeticus* tyrosinase expression. **(A)** Chemical pathway of eumelanin production from L-tyrosine. The first step involves the hydroxylation of L-tyrosine to L-DOPA which is catalysed by tyrosinase – here acting as an monophenol monooxygenase. This step is then followed by the catalysis of L-DOPA to dopaquinone, which is catalysed by the diphenolase activity of Tyr1. The remaining steps in the pathway occur spontaneously in the presence of oxygen, leading to the generation of eumelanin. **(B)** Genetic construct maps for two *K. rhaeticus* tyrosinase expression strains: *K. rhaeticus* _*p*_*tyr1* and *K. rhaeticus tyr1*. Both constructs use the same constitutive promoter (*pJ23104*), RBS (*B0034*) and terminator (*L3S1P00*). *K. rhaeticus* _*p*_*tyr1* uses a plasmid with a pBBR1 origin of replication and a chloramphenicol resistance cassette. **(C)** A two-step process for eumelanin production from *K. rhaeticus* grown in shaking conditions. Strains are grown in HS-glucose media, washed and resuspended with PBS to remove spent media before being mixed with melanin development buffer. **(D)** Tyr1 producing strains are assayed for eumelanin production. Eumelanin production per cell was determined by measuring OD_405_ over 12 hours, divided by the initial OD_600_ of each well at time point 0. **(E)** Initial reaction rate per cell was determined by measuring the gradient of eumelanin accumulation per cell from 50-170 minutes after the start of measurement. Error bars are the standard deviation of three biological replicates. **(F)** Optical microscopy images of *K. rhaeticus* _*p*_*tyr1* before (mel-) and after melanin development (mel+). A zoom-in example of a single cell is shown with a cyan outline.

Melanin synthesis by Tyr1 is sensitive to pH — only occurring readily at pH values above 7^42^. This conflicts with the growth of *K. rhaeticus*, which as an acetic acid bacteria, acidifies its culture media during growth by the production of organic acids such as gluconic and acetic acid^43,44^. Indeed, we found that *K. rhaeticus* _*p*_*tyr1* pellicles grown in Hestrin-Schramm glucose (HS-glucose) media buffered to pH 5.7 and with the necessary substrate and co-factors for eumelanin production (0.5 g/L L-tyrosine and 10 μM CuSO_4_), displayed no pigmentation during growth (**Extended data: Fig. 1a**)^45^. We measured the acidification of the growth media after pellicle production, which demonstrated that the culture pH had lowered to below pH 4, even when initial media pH was buffered higher to pH 7 **(Extended data: Fig. 1b**). These results suggested we would need to separate pellicle growth from eumelanin production.

We therefore decided to employ a two-step approach to produce melanin from *K. rhaeticus*. Step one would involve growing Tyr1-expressing *K. rhaeticus* under normal growth conditions and step two would involve removing the spent culture media and replacing it with a buffered solution with the reagents required for melanin synthesis (**Fig. 1c**). For the buffered solution, which we refer to as *melanin development buffer*, we chose to use phosphate-buffered saline (PBS), buffered to pH 7.4, containing 0.5 g/L L-tyrosine and 10 μM CuSO_4_. We then tested this approach with both of our Tyr1-expressing strains. To enable quantification of eumelanin production, we assayed *K. rhaeticus* cells that had grown in shaking conditions with cellulase added to the media, which prevents pellicle formation. We measured eumelanin production in the melanin development buffer at OD_405_ over 12 hours (**Fig. 1d**).

Both Tyr1-expressing *K. rhaeticus* strains were able to produce eumelanin in the development buffer. The melanin production rate per cell was higher for the integrated tyrosinase strain *K. rhaeticus tyr1* (0.48 ± 0.03 OD_405_/OD_600_/h) vs. the plasmid-based *K. rhaeticus* _*p*_*tyr1* (0.35 ± 0.02 OD_405_/OD_600_/h) **(Fig. 1e**). We also used this same experimental approach to assay the effect on melanin production of changing the pH, salt concentration, oxidation state (II) metal ion and copper (II) concentration of the melanin development buffer (**Extended data: Fig. 2a-h**). Interestingly, we found alkaline buffer conditions (>pH 8) led to a more rapid production of eumelanin than neutral conditions. However, the production rate in alkaline buffer conditions also slowed rapidly, leading to overall lower melanin accumulation than at pH 8. This result conflicted with previous *in vitro* studies on Tyr1 that suggested an optimal pH value of 7^42^. The same study did state, however, that L-DOPA may spontaneously convert to dopachrome at pH values above 7.5. The *in vivo* result seen here may reflect a spontaneous conversion of built-up L-DOPA inside the cells, which may suggest Tyr1 has some level of monophenolase activity (L-tyrosine to L-DOPA) during the *K. rhaeticus* growth stage, despite a lack of visible melanin production.

**Figure 2.**
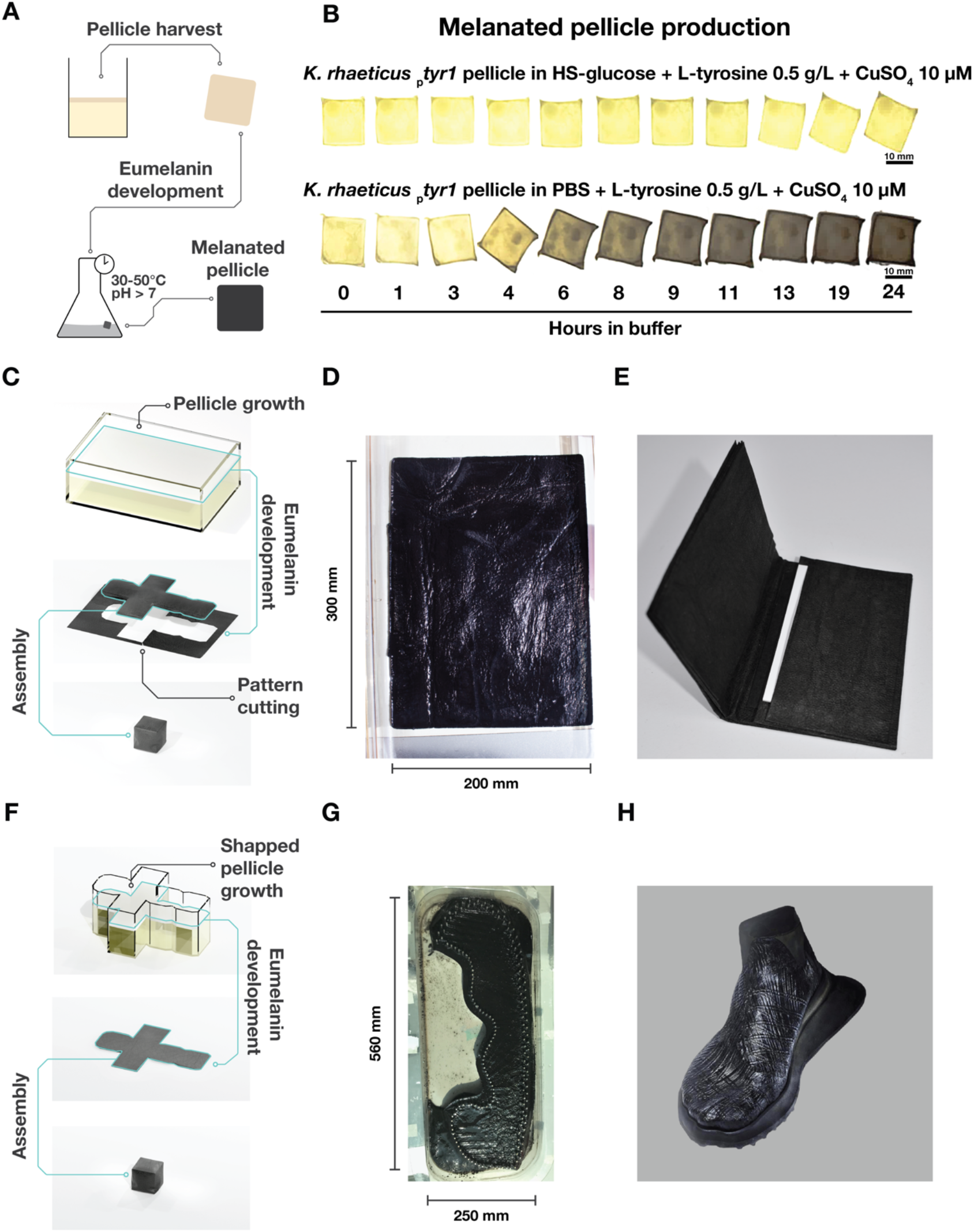
Using *tyr1* expressing *K. rhaeticus* to produce melanated BC. **(A)** The production process for melanated bacterial cellulose involves two steps. Tyrosinase-expressing *K. rhaeticus* are grown in static conditions to produce a pellicle. Once grown, this is harvested and placed in development buffer and incubated with agitation between 30°C and 50°C until the material reaches the desired shade. **(B)** Images show a time-lapse of the progression of eumelanin accumulation over 24 hours for a *K. rhaeticus* _*p*_*tyr1* pellicle placed in either development buffer or HS-glucose media. **C)** Pellicle production can be conducted in standardised containers to produce sheets of BC from which pattern pieces can be cut out and assembled. **(D)** A *K. rhaeticus tyr1* pellicle, grown in a 300×200 mm container, after eumelanin development step. **(E)** A finalised wallet prototype, cut and assembled from two pressed and dried melanated BC sheets. **(F)** Pellicle production can also occur in shaped containers, producing BC pre-shaped to the 2D pattern of the final pattern piece. **(G)** A *K. rhaeticus* _*p*_*tyr1* pellicle grown in a shaped container, after eumelanin development. **(H)** A finalised shoe upper prototype produced from a melanated shaped BC sheet that has been wrapped around a foot shaped last and allowed to dry.

We then looked at how *K. rhaeticus* cells had changed after exposure to the melanin development buffer. Using light microscopy, we found that *K. rhaeticus* _*p*_*tyr1* cells exposed to the development buffer appeared visibly darker suggesting eumelanin production may be occurring intracellularly (**Fig. 1f**). This was as expected, given that the Tyr1 protein did not contain secretion or translocation tags. However, as the onset of eumelanin production requires the cells be immersed in a neutral pH buffer this also suggests that the cytoplasmic pH of *K. rhaeticus* may also become acidic during growth. Indeed, other acetic acid bacteria show adaptations suggestive of acidic internal conditions^43,46^.

### Pigmenting BC through *K. rhaeticus* eumelanin production

Having shown eumelanin production from *K. rhaeticus* cells expressing *tyr1*, we next wanted to demonstrate that eumelanin production could effectively pigment bacterial cellulose (BC). To do so, we applied the same two-step process to a *K. rhaeticus* _*p*_*tyr1* static culture that had grown a pellicle (**Fig. 2a**). Following 24 hours of shaking incubation at 30°C in the development buffer, the pellicle changed its appearance from a pale yellow to a brownish black, demonstrating eumelanin pigmentation of BC (**Fig. 2b**). Additionally, by reducing L-tyrosine concentration in the development buffer we could slow the rate of melanin production (**Extended Fig. 3a**), allowing us to vary how pigmented a BC pellicle becomes and thus generate material in a range of brown shades (**Extended Fig. 3b**). We also found that including 0.5 g/L L-tyrosine in both culture medium as well as the melanin development buffer led to the darkest pellicles, presumably as this allows L-tyrosine levels and eumelanin precursors to build-up in the cells during growth. Finally, crucial to the use of melanated BC outside of laboratory contexts is that the pigment persists through sterilisation. We found that both high-pressure steam and ethanol sterilisation worked well to preserve pigmentation (**Extended Fig. 3c**). As expected, sterilisation by oxidising compounds, such as sodium hypochlorite bleach, led to a rapid loss of melanin pigmentation.

**Figure 3.**
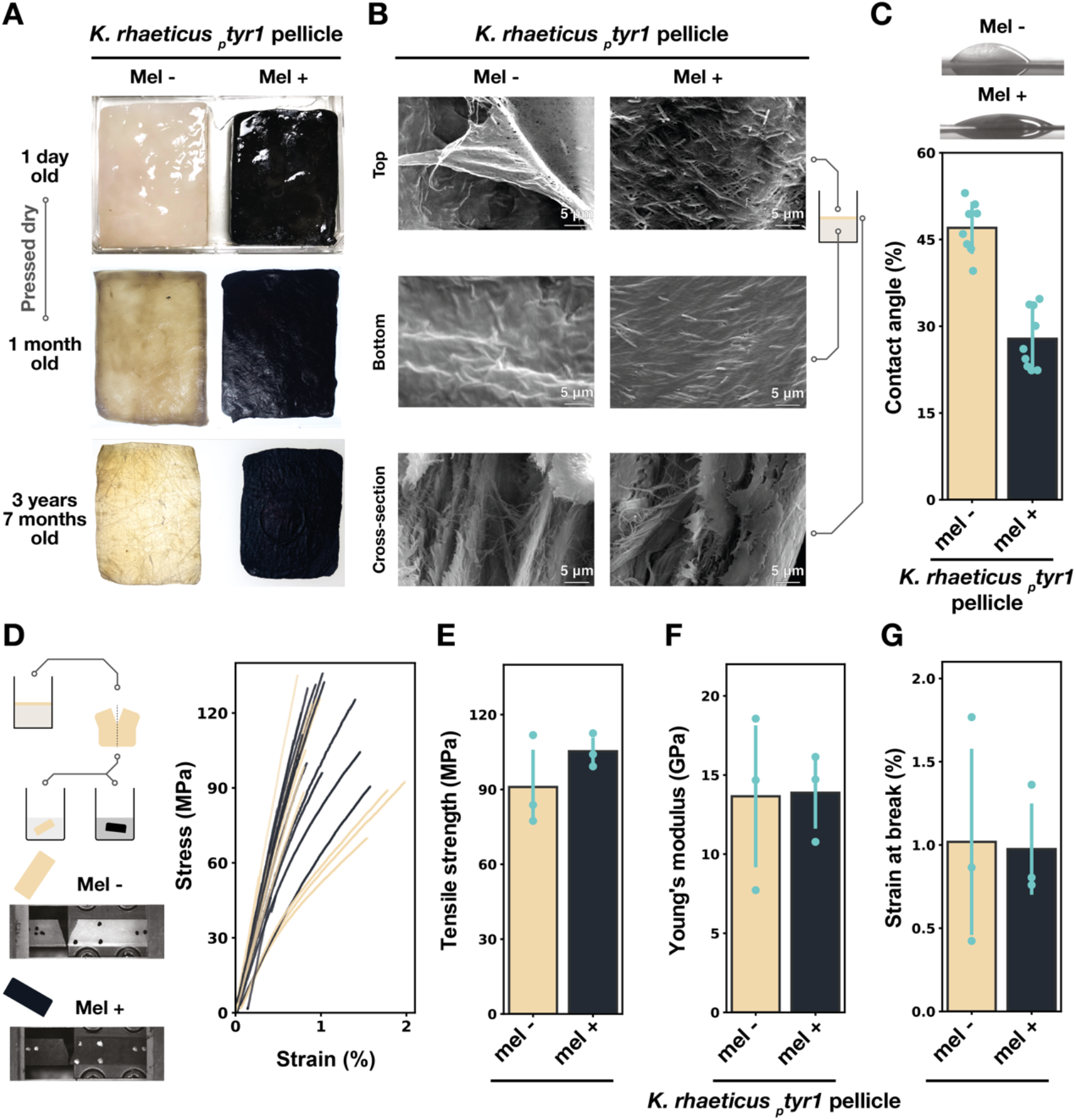
Properties of melanated bacterial cellulose. **(A)** A melanated (mel+) and unmelanated (mel-) swatch, from the same original *K. rhaeticus* _*p*_*tyr1* pellicle. These swatches had been dried and used as demonstration pieces. **(B)** Scanning electron microscopy of mel+ and mel-*K. rhaeticus* _*p*_*tyr1* bacterial cellulose. The top and bottom surfaces pertain to the air-facing and media-facing pellicle surfaces respectively. **(C)** The sessile drop method was used to measure the contact angle on *K. rhaeticus* _*p*_*tyr1* mel-(yellow) and mel+ (black) pellicles. An unpaired T-test result gave a p value < 0.005 and error bars represent standard deviation from 8 mel- and 9 mel+ drop measurements. Representative water drop shapes for mel+ and mel-pellicles are shown above the graph. **(D)** Comparative Tensile tests of melanated and unmelanated BC nanofibril networks were made using pressed networks prepared from halves of the same *K. rhaeticus* _*p*_*tyr1* pellicle. Representative images of BC breaks are given for mel- and mel+ BC as well as stress-strain curves of the technical repeats from 3 biological replicates of mel+ and mel-pellicles. **(E-G)** Tensile strength, Young’s modulus and Strain at Break for mel+ and mel-pellicles — the P values from paired T-tests between mel+ and mel- BC were 0.17, 0.92, 0.85 respectively. Error bars show standard deviation from 3 biological replicates and each biological replicate is the average of 3 or more technical replicates.

Due to high yields of material being produced from simple static growth cultures, microbial production of BC is very amenable to scale-up, enabling the amounts of BC required to build real products to be achieved with minimal infrastructure investment. This has made BC attractive to producers at both industrial and *cottage-industry* scale, especially as a vegan alternative to leather for use in clothing and accessories. With this in mind, we wanted to demonstrate that we could scale growth of Tyr1-expressing *K. rhaeticus* to produce functionally useful quantities of pigmented BC. For this we looked at two approaches to BC production. In our first approach, we sought to produce a standardised sheet of melanated BC, from which a textile pattern (*i.e*. template) could be cut and assembled (**Fig. 2c**). To aid in the large-scale growth of BC, we switched to growth media containing coconut water, 1% ethanol and 1% acetic acid. This media is used in industrial environments to grow *K. rhaeticus* and maximise BC production^47^. We grew *K. rhaeticus tyr1* in a 300×200 mm tray and after 10 days growth we harvested the pellicle and let it undergo eumelanin production until it had taken on a deep black colour (**Fig. 2d**). The melanated BC sheet was then sterilised by autoclave, pressed flat and dried. The BC sheet retained its colour throughout this process, and the dried sheet had a paper-like but flexible feel (**Supplementary video 1**). A wallet pattern was then cut from two of these melanated sheets, and the pattern pieces sewn together with thread to make a functioning melanated BC wallet (**Fig. 2e**).

In our second approach we took advantage of how pellicle growth follows the air:water interface and grows in the same shape as the culturing vessel (**Fig. 2f**). Using *K. rhaeticus* _*p*_*tyr1* we grew a pellicle in a bespoke culture vessel, in the shape of a shoe-upper pattern piece. This culture vessel contained a loom-like apparatus holding a network of strung Lyocell (TENCEL) threads, located at the air:water interface to allow these threads to be incorporated into the growing *K. rhaeticus* _*p*_*tyr1* pellicle. After 14 days growth, the final pellicle and apparatus were removed from the culture media and placed into development buffer. After 48 hours of gentle shaking at 30°C the pellicle had taken on a deep black colour (**Fig. 2f**). The pellicle was then sterilised by ethanol bath and soaked in a 5% glycerol solution, before being removed from the apparatus and wrapped around an epoxy shoe last (*i.e*. a foot-shaped mold) and allowed to dry **(Fig. 2g**).

Both products demonstrate that our engineered strains can grow and self-pigment at scales large enough to produce viable prototype fashion pieces. They also demonstrate the positive outcomes of collaboration between scientists and designers in the pursuit of creating new engineered living materials. As the first users of new biomaterial-based textiles, designers play a key role in demonstrating and publicising the features of a new material and can give constructive feedback to scientists on any limitations and how the materials could be improved, particularly in order for their end-of-life to become more sustainable.

### Characterisation of melanated cellulose

A swatch of melanated cellulase produced by *K. rhaeticus* _*p*_*tyr1* was actively used as a demo piece for 42 months and maintained its pigmentation throughout (**Fig. 3a**), demonstrating that the colour was resilient over time. As well as colour, we were curious to know how eumelanin production may have impacted the other material properties of bacterial cellulose. To investigate this, we first checked to see if eumelanin had altered the BC surface using scanning electron microscopy (SEM). We compared the top and bottom surfaces, as well as the cross-sections, of melanated and unmelanated *K. rhaeticus* _*p*_*tyr1* pellicles **(Fig. 3b**). The SEM images do not indicate any major structural differences in the surface architecture between melanated and unmelanated cellulose. To further study the surface material properties of melanated cellulose, we conducted wettability testing using the static sessile drop method (**Fig. 3c**). Using pellicles grown from *K. rhaeticus* _*p*_*tyr1*, we observed that the melanated pellicle had increased surface wettability, with an average contact angle of 28° compared to 47° for the unmelanated pellicle.

One of the most attractive features of bacterial cellulose for industry is its high tensile strength, therefore, it is important to know if melanation interferes with or enhances the strength of the BC nanofibril network. We carried out tensile testing using both melanated and unmelanated pellicles. For consistency, we prepared a paired set of BC samples, by splitting each grown pellicle in half, developing eumelanin in only one half of each pellicle and pressing both halves into a consolidated network of BC nanofibrils. (**Fig. 3d**). The average tensile strength values were 91 MPa and 105 MPa for unmelanated and melanated pellicles respectively (**Fig. 3e**). For the Young’s modulus, the values were 13.7 GPa and 13.9 GPa for unmelanated and melanated pellicles respectively (**Fig. 3f**). Strain at break was 1.02% and 0.98 % for Unmelanated and melanated samples respectively (**Fig. 3g**). The material properties of all samples tested fell within the expected ranges of 70-300 MPa for BC tensile strength and 5-17 GPa for Young’s modulus^30^. A paired T-test showed that the BC nanofibril networks prepared from melanated and unmelanated pellicles did not possess significant statistical differences in tensile material properties.

### Patterning eumelanin output

Beyond colouring, textile processing can also involve patterning a textile. To showcase the capability of genetically engineered self-dyeing BC textiles, we set out to establish spatial control of gene expression. We reasoned that patterning of gene expression within a growing material may create durable pigmentation that would extend vertically through the material, a feature not seen in synthetic leathers that rely on printed on surface patterns that are protected with a finish. We determined that the most flexible and precise way to pattern gene expression would be through engineered optogenetics and proposed a procedure for making patterned BC with light (**Fig. 4a**). To engineer a light-sensitive strain of *K. rhaeticus*, we chose to implement the blue-light sensitive T7-RNA polymerase (Opto-T7RNAP) system originally designed for use in *E. coli* by Baumschlager *et al*. (**Fig. 4b**)^48^. We surmised that the Opto-T7RNAP system would be one of simplest optogenetic systems to implement in a non-model organism like *K. rhaeticus*^49^. This view was based on the orthogonality of T7-RNA polymerase transcription; the lack of membrane-bound light-sensing components and the use of ubiquitous flavin adenosine dinucleotide (FAD) as a chromophore, which eliminates the need for heterologous chromophore biosynthesis genes.

**Figure 4.**
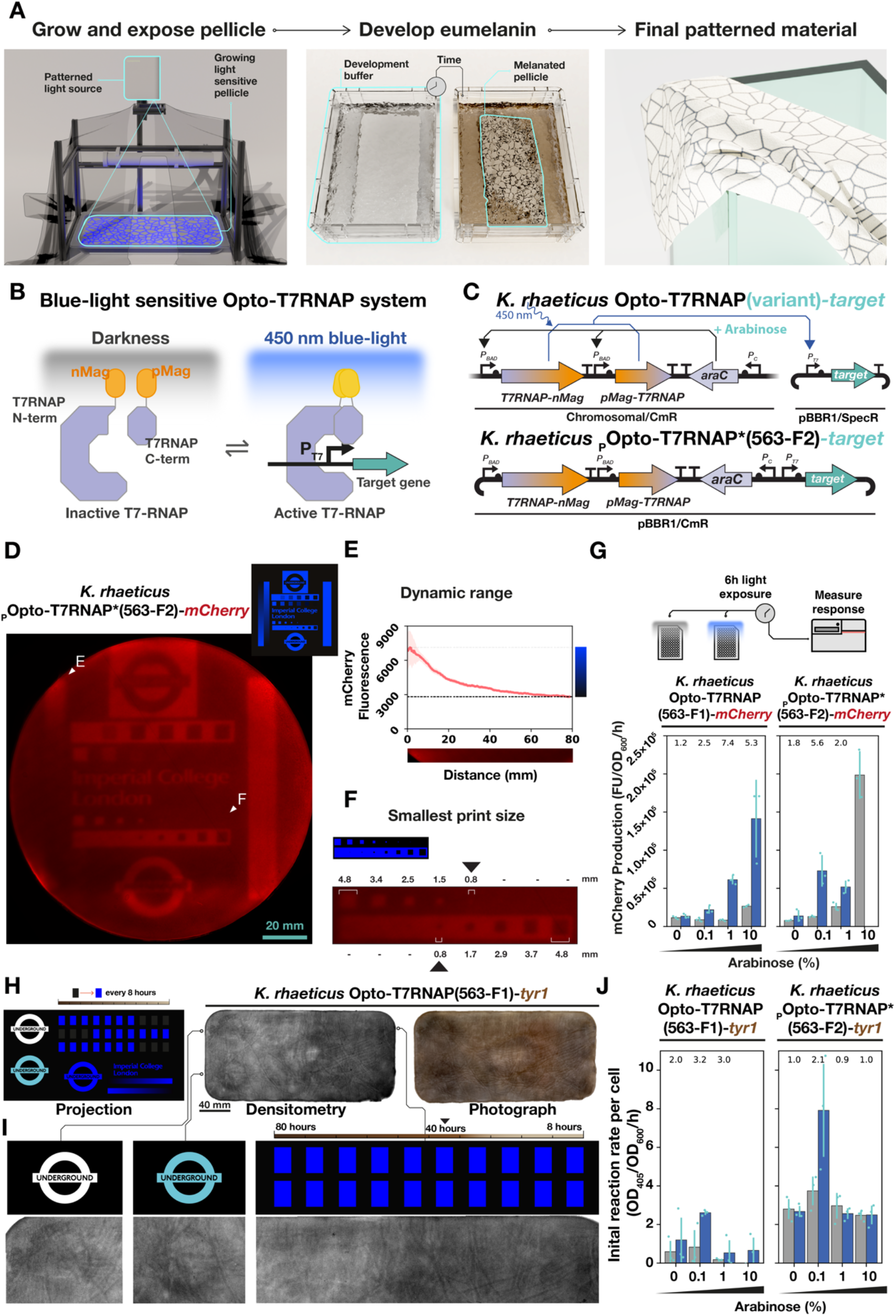
Functional optogenetics in *K. rhaeticus*. **A)** Proposed procedure to make patterned bacterial cellulose through optogenetics. **B)** The Opto-T7RNAP system uses a split T7 RNA polymerase, with each half fused to a photo-sensitive magnet protein (orange). Under blue light (∼450 nm), the magnet proteins form a dimer (yellow) that initiates transcription from a P_T7_ promoter. **C)** Genetic construct maps for two arrangements of *K. rhaeticus* optogenetic strains. In both strains, AraC is transcribed constitutively. Addition of arabinose activates the expression of the split T7 RNAP genes from both _p_BAD promoters. **D)** Red fluorescence scan of the top surface of a blue light exposed *K. rhaeticus* _P_Opto-T7RNAP*(563-F2)*-mCherry* pellicle. Pellicle displayed has a diameter of 150 mm. Graphic on the top right shows image projected onto the pellicle during growth. **E)** The left of the projected image contained a gradated strip, from minimum to maximum light let through. Data show intensity of red fluorescence seen in the pellicle against this gradated strip. Pink area shows the standard deviation of pixel intensity at each horizontal slice. **F)** The smallest projected mark on the exposed pellicle. **G)** Characterisation of optogenetics constructs with *mCherry* target gene under differing arabinose % (w/v) concentration. Bars (blue = exposed, grey = unexposed) show average increase in red fluorescence after 6 hours normalised by OD_600_. Error bars show standard deviation of 3 biological replicates placed on the same microtiter plate. Fold difference between exposed and unexposed cells is shown at top, except in instances of poor cell growth. **H)** Comparison between projection image and the resulting *K. rhaeticus* Opto-T7RNAP(563-F1)-*tyr1* pellicle after eumelanin development. Blue rectangles are added to the top right of the projection image every 8 hours to test required exposure time -progressing from 1 rectangle to 10 rectangles per line over 80 hours. A densitometry scan and photograph of the top surface of the pellicle is shown. Pellicle was 300 × 170 mm in size. **I)** Zoomed-in sections of a densitometry scan of the *K. rhaeticus* Opto-T7RNAP(563-F1)-*tyr1* pellicle compared to the projection. The black triangle points to the 6th rectangle, which equates to a 40-hour required exposure time. **J)** Characterisation of optogenetic constructs with *tyr1* target gene under differing arabinose induction. Data show the average and standard deviation of three biological replicates of initial (0 – 100 mins) reaction rate of eumelanin production measured at OD_405_, normalised to the number of cells at OD_600_ at time point 0.

To implement the Opto-T7RNAP system in *K. rhaeticus* we tested two arrangements of the necessary DNA parts and selected the variant with the highest fold-change between light and dark states in Baumschlager *et al*. -Opto-T7RNAP*(563-F2) -as a basis of a plasmid-based version, where the light-sensitive T7RNAP genes and light-regulated target gene were placed on the same plasmid (**Extended data: Fig. 4a**). In an alternative arrangement, to test variants of the Opto-T7RNAP system produced in Baumschlager *et al*., we integrated the light-sensitive T7RNAP variant genes into the *K. rhaeticus* chromosome. These *K. rhaeticus* T7RNAP variant strains were then transformed with a separate plasmid encoding the target gene (**Fig. 4c**). In both arrangements, the two Opto-T7RNAP light-sensitive genes were regulated by the P_BAD_ promoter, which had been previously shown to function in *K. rhaeticus* when induced with arabinose at a concentration of 2% (w/v)^50^. The arabinose regulator gene, *araC*, was placed downstream of the integrated two Opto-T7RNAP light sensitive genes in both the chromosomal and plasmid arrangements.

We then tested whether we could pattern gene expression in *K. rhaeticus* using the Opto-T7RNAP system. To simplify this process, we started with our target gene being a fluorescent reporter gene that produces the red fluorescent protein (RFP) mCherry. We constructed a projection device that could project an image onto the surface of culture liquid as it grows a pellicle (**Extended data: Fig. 4b**). We inoculated *K. rhaeticus* _p_Opto-T7RNAP*(563-F2)-*mCherry* in the culturing vessel of this device, and once a thin pellicle had formed, we exposed this nascent pellicle to a projected image for 72 hours. The harvested pellicle showed successful pattering of *mCherry* expression (**Fig. 4d**). The pellicle showed a 2.54x fold change in fluorescence between the least and most exposed section of the pellicle (**Fig. 4e**). Importantly, we also found the smallest region we could visually pattern was 0.8 mm^2^, which gave us a working estimate of the resolution of the approach (**Fig. 4f**).

We then set out to understand the optimal level of light-sensor expression in *K. rhaeticus* to maximise the dynamic range between light and dark states. We studied this by measuring blue-light responses from the *K. rhaeticus* Opto-T7RANP variants exposed to a range of arabinose concentrations when grown with shaking conditions in parallel in microtitre plates (**Extended data: Fig. 4c**). After evaluating the *mCherry* expression response from exposure to blue-light for 6 hours, we found that only one of the integrated variants -*K. rhaeticus* Opto-T7RNAP(563-F1)-*mcherry* -showed an *mCherry* expression level above that of *K. rhaeticus* _*p*_*T7-mCherry*, the strain containing just the target plasmid –_p_*T7-mCherry* – with no T7-RNA polymerase **(Fig. 4g**). The maximal fold change from *K. rhaeticus* Opto-T7RNAP(563-F1)-*mCherry* was at 1% (w/v) arabinose and gave a 7.4x fold change in mCherry production. For the plasmid-based *K. rhaeticus* _p_Opto-T7RNAP*(563-F2)-*mCherry*, 0.1% arabinose was found to be the preferred condition for dark to light switch behaviour, whereas 10% arabinose overloaded the circuit and gave high mCherry production in the dark, and no detectable fluorescence at all when given light.

Having demonstrated we could pattern gene expression with the Opto-T7RNAP system and light projection, we next moved to patterning eumelanin accumulation in a pellicle. Having successfully used *K. rhaeticus* _p_Opto-T7RNAP*(563-F2)-*mCherry* to pattern mCherry RFP expression, we decided to create an alternative version of this strain where the *mCherry* coding sequence was directly replaced with the *tyr1* coding sequence, creating *K. rhaeticus* _p_Opto-T7RNAP*(563-F2)-*tyr1*. However, when tested in parallel with the *mCherry* version, we found the *tyr1* version had such a high level of background eumelanin production, that it obscured visible patterning of eumelanin accumulation (**Extended data: Fig. 4d)**. We therefore switched to the other variant that had also shown an appreciable blue-light response in the *mCherry* testing -*K. rhaeticus* Opto-T7RNAP(563-F1)-*tyr1* – to further test if we could pattern eumelanin accumulation.

Ahead of testing we designed a dynamic image to be projected and a new projection set up with a commercial projector to allow us to change the image during the exposure and thus measure the required exposure times to generate an appreciable eumelanin response **(Extended data: Fig 4e**). Using *K. rhaeticus* Opto-T7RNAP(563-F1)-*tyr1*, we grew a BC pellicle in this device, and once grown, we exposed the pellicle to an 80 hour projection (**Fig. 4h**). We then took the exposed pellicle, placed it into development buffer and incubated it at 30°C for 48 hours by which point we could observe rough patterning of eumelanin accumulation (**Fig. 4i**). Unfortunately, while the pellicle shows some evidence of patterning in places, a high degree of background eumelanin accumulation also made this patterning attempt hard to decipher. We could, however, determine from the dynamic patterning that at least 40 hours was required to observe visible eumelanin accumulation when it is produced from *K. rhaeticus* Opto-T7RNAP(563-F1)-*tyr1*.

Finally, we assessed how the other Opto-T7RNAP variants behaved with *tyr1* as their target gene. We used a similar parallel microtitre plate approach as before but now measuring the accumulation of eumelanin per cell over time at OD_405_ (**Fig. 4j**). This revealed that *K. rhaeticus* _p_Opto-T7RNAP*(563-F2)-*tyr1* required 0.1% arabinose to work correctly, as it had when *mCherry* was the target (**Fig. 4g**). However, now in this condition and all others tested, a high eumelanin production rate is seen even in the absence of light induction. This finding agrees with the high background pigmentation seen in the test pellicle. Among the strains with chromosomally-integrated DNA, *K. rhaeticus* Opto-T7RNAP(563-F1)-*tyr1* showed the highest eumelanin production rate in response to blue light, again requiring 0.1% arabinose to tune this. The fold change between the light and dark states for this strain was lower than that for two other variants tested, which both showed fold changes greater than 10x (**Extended data: Fig. 4F**). However, these strains are not ideal for pigmentation, as they have much lower eumelanin production rates. Overall, while we can show the control of eumelanin production can be regulated with blue-light using the Opto-T7RNAP system, accurate patterning of eumelanin accumulation in a pellicle remains to be optimised.

Our main limitations to patterning eumelanin accumulation using Opto-T7RNAP were high levels of background pigmentation and a constrained fold-change in response to blue light. These two factors severely reduced the dynamic range of the system. Other factors that could have also reduced pattern definition in the pellicle include Tyr1 enzyme or L-DOPA leaking from cells (e.g. via cell lysis), or *tyr1* expression leading to reduced cell growth and density in blue-light exposed regions. Most of these factors could be addressed by decreasing background *tyr1* expression from the target plasmid, but the exact source of this background expression is currently unknown. Optimisation of the arrangement of the Opto-T7RNAP genes in *K. rhaeticus* will hopefully allow us to approach the dynamic range of the system seen in *E. coli*^48^. We anticipate that improving the performance of the Opto-T7RNAP system in *K. rhaeticus*, and developing alternative target genes, will yield more advanced BC biomaterials in the near future.

## Discussion

To the best of our knowledge this project is the first to employ genetic engineering in a material-producing bacteria for the purpose of creating pigmented products. We have demonstrated that tyrosinase expression from *K. rhaeticus* is proficient to produce highly pigmented BC; that growth of engineered *K. rhaeticus* can be scaled to produce useful quantities of pigmented BC and that this pigmentation is stable. This work demonstrates the value of using genetic engineering to design and construct strains intended to grow materials with desired properties; in this case with a chosen colour grown into the material, rather than having to be added to it later by an industrial chemical dyeing process.

Our work represents only a first step in the development of melanated BC. The use of the two-step process for eumelanin production increases the amount of water required for melanated BC and this may restrict its sustainability. Potentially this additional water usage could be reduced through the reuse and recycling of development buffer, or the addition of the buffer components directly to the growth media after pellicle production. However, the most effective approach may be to adapt eumelanin production to occur in the acidic conditions generated by *K. rhaeticus*. Such an approach will require an evaluation of the eumelanin chemical pathway to identify the sources of sensitivity to low pH, and a survey of natural melanin biosynthesis mechanisms to identify acid-tolerant tyrosinases and/or acid tolerant eumelanin biosynthesis.

We are confident that the production of melanated BC can be scaled further in an industrial context. While we’ve demonstrated that Tyr1 functions inside *K. rhaeticus*, many other *Komagataeibacter* species are used in industrial BC production and the culturing conditions to maximise yields from each species are quite varied. Therefore, to increase the flexibility of our work, the genetic engineering of *tyr1* expression should next be demonstrated in other *Komagataeibacter* species. At the industrial level, it will be important that melanated BC is subject to the testing of industry standards for colour fastness, such as crock testing and resistance to fading in UV and visible light. Additionally, the sustainability credentials of melanated BC should also be assessed through full life-cycle analysis. Melanated BC will also face the challenges that already limit the use of BC as a textile. Chiefly, the high hydrophilicity of BC mandates significant waterproofing during the textile processing stage. Potentially, however, this process could also be reconsidered by genetic engineering of *Komagataeibacter*, either to alter the grown BC structure, or through the biosynthesis of a layer of hydrophobic compounds. Additionally, the sterilisation of BC materials, especially in the case of high-pressure steam sterilisation, carries significant energy requirements that will need to be considered.

Finally, there are additional avenues to explore the genetic engineering approach, demonstrated here, to produce other pigment molecules from *K. rhaeticus*. The L-DOPA produced by tyrosinase can, in the presence of cysteine, be shifted to the formation of the red pigment, pheomelanin, and further work here would expand the possible shade range of melanated BC materials^51^. The biosynthesis of other insoluble pigment molecules could also be pursued. The most obvious being the production of indigoid dyes, such as indigo and tyrian purple, whose biosynthesis has already been demonstrated in *E. coli*^52,53^. Indeed, the engineered biosynthesis of only a small range of pigments could deliver a supernumerary range of BC shades, due to the potential of mixing pigment production at the genetic level and through the co-culturing of different pigment producing strains.

## Methods

### *K. rhaeticus* culture conditions and culturing approaches

Two culture media were used in this study to culture *K. rhaeticus* Hestrin–Schramm (HS-glucose) media (2% glucose, 10 g/l yeast extract, 10 g/l peptone, 2.7 g/l Na2HPO4 and 1.3 g/l citric acid, pH 5.6–5.8) and coconut water media (coconut water (Vita Coco), 0.05% (v/v) acetic acid). Coconut water media was sterilised by filtration, except in situations where over 1L were required. In those situations, media supplements were sterilised separately and combined with coconut water, which had been opened and decanted out with aseptic technique, in the culturing container.

When *K. rhaeticus* was cultured on solid media, HS-glucose media was always used and supplemented with 1.5% agar. *K. rhaeticus* liquid cultures fell into two separate approaches: shaking cultures and stationary cultures. In shaking cultures, the media in use was supplemented with 2% (v/v) cellulase (Sigma-Aldrich: C2730) to allow for turbid growth without clumping. In stationary culture, where the goal is pellicle formation, media would be supplemented with 1% (v/v) ethanol to enhance pellicle production. In both approaches, where antibiotics were required for plasmid maintenance, media was supplemented with 340 μg/ml chloramphenicol or 200 μg/ml spectinomycin.

To facilitate consistency when inoculating multiple pellicles, *K. rhaeticus* cells would be grown in shaking growth conditions until turbid, normalised in OD_600_ across samples, pelleted by centrifugation and washed in the subsequent media to remove cellulase. The washed cells were used as a pre-culture and added, at a ratio of 1:25, into the culturing container and left in stationary conditions at 30°C to form pellicles. In the case of forming large pellicles consistently (>25 cm^2^), a *glycerol aliquot* approach was used. The *K. rhaeticus* strain of interest would be grown, shaking at 30°C in 100 ml of HS-glucose media until it reached an OD_600_ ∼0.6-1. At this point, the cells would be pelleted by centrifugation, washed in HS-glucose media, before being pelleted again and resuspended in 10 ml of HS-glucose media containing 25 % glycerol. The resuspended cells would be separated into 1 ml aliquots and stored at −80°C until use. When used, an aliquot would be thawed and added to the media in the final culturing container.

### Molecular biology and strain construction

DNA parts and plasmids used in this study are listed in the supplementary materials. *E. coli* Turbo (NEB) cells were used for plasmid construction. The *tyr1* DNA sequence was ordered from Twist Bioscience, with compatible 3’ and 5’ overhangs for entry into the KTK via Golden Gate Cloning. Constitutive tyrosinase constructs were built using the KTK. The procedures and protocols of working with the KTK are described in Goosens *et al*. Plasmids containing the various versions of the Opto-T7RNAP system were kindly sent to us by Armin Baumschlager and Mustafa Khammash from ETH Zürich. Due to the presence of multiple KTK-incompatible restriction sites in the T7-Opto coding sequences, Gibson cloning was used to build both the _p_Opto-T7RNAP*(563-F2)*-target* plasmid and the 5 _p_Opto-T7RNAP suicide plasmids for genomic integration. The primers for Gibson cloning are listed in the supplementary materials.

*K. rhaeticus* electrocompetent cells were prepared as in Florea *et al*^23^. *K. rhaeticus* cells were transformed using electroporation and selected for on HS-glucose agar plates containing either 340 μg/ml chloramphenicol or 500 μg/ml spectinomycin, depending on the plasmid selection marker in use. Note, here a higher concentration of spectinomycin is used than during normal culturing. Genetic constructs that were integrated into the chromosome of *K. rhaeticus*, were done so by homologous recombination using a pUC19 suicide plasmid, as described in Goosens *et al*.

### Melanated pellicle production

Melanated pellicles were produced using a two-step approach. First a *tyr1* expression strain would be inoculated into a sterile culture container. Typically, 24 well deep well plates (Axygen) were used to make small pellicles. Each well contained 5 mL of growth media and was inoculated at a 1:25 ratio with pre-culture. Growth media was enriched with 0.5 g/L L-tyrosine and 10 μM CuSO_4_ to promote the highest eumelanin production. Once the pellicles had reached the desired thickness, they were harvested, placed in a bath of sterile dH_2_O and washed for 1 minute by gently shaking by hand. The washed pellicles are then passed into a bath of eumelanin development buffer. A large ratio of buffer to pellicle was used, *i.e*. one pellicle in 25 mL of buffer in a 50 mL falcon tube, this was to prevent the overwhelming of the buffer by remaining acid in the pellicle. The pellicle would be allowed to produce eumelanin at >30°C in shaking conditions over 24 hours.

### Large melanated pellicle production

To produce the melanated pellicle used to make the wallet, a 200×300 Eurobox container was sterilised and filled with 3 L of coconut water media supplemented with 0.5 g/L L-tyrosine, 10 μM CuSO_4_ and 1% ethanol. The media was inoculated with a 1 ml *K. rhaeticus tyr1* glycerol aliquot and covered in paper towel before being placed into a stationary incubator set to 30°C. After 10 days growth, the pellicle was harvested, washed briefly in dH_2_O before being placed in a 300×400 mm Eurobox containing 2 L of concentrated eumelanin development buffer (10x PBS). The development container was then placed into a shaking incubator set to 45°C and allowed to produce eumelanin over 2 days, at which point the cellulose had become completely black. The melanated cellulose was then washed again to remove excess eumelanin development buffer before being autoclaved. To make the material pliable after drying, the cellulose sheet was left in a 5% glycerol solution, before being pressed and dried.

To produce the melanated pellicle used to make the shoe, a custom shaped vessel, containing an apparatus that held a network of tight strung yarn, was sterilised and filled with 2 L of coconut water media supplemented with 0.5 g/L L-tyrosine, 10 μM CuSO_4,_ 340 μg/mL chloramphenicol and 1% ethanol. The media was inoculated with a ∼500 mL pre-cultured *K. rhaeticus* _*p*_*tyr1* pellicle. The culture was left to grow at room temperature in stationary conditions, until a thin pellicle had formed. At this point, fresh coconut water media supplemented with 0.5 g/L L-tyrosine, 10 μM CuSO_4,_ 340 μg/mL chloramphenicol and 1% ethanol was added, to raise the pellicle to just below the level of the tensed yarn. After two weeks’ growth, the media was drained and replaced with concentrated eumelanin development buffer (10x PBS). The full container was placed into a shaking incubator set to 30 RPM, and developed at 30°C for 1 day, at which point the pellicle had become completely black. The vessel was then drained of eumelanin development buffer, replaced with 70 % ethanol and left overnight to sterilise. The ethanol was replaced with a 5 % glycerol solution before the melanated cellulose was removed from the apparatus and wrapped around a shoe shaped mold to dry.

### Eumelanin production assay

The eumelanin production assay uses a 384 square-well microtiter plate as a reaction plate. An OT-2 liquid handling robot (Opentrons) was used to prepare these reaction plates for the assay. Development buffer was placed into a deep well plate, from which 40 μl was transferred to each well in the reaction plate using an 8-channel 300 μL OT-2 Gen2 pipette. The reaction plate was kept at 4°C to slow eumelanin production during plate preparation using the OT-2 Thermo-module. Cells potentially containing tyrosinase were placed into a 96 well plate. Cells were mixed in one round of aspiration using an 8-channel 20 μL OT-2 Gen2 pipette before 10 μL of cells were transferred into each well of the 384 well plate. Once full, the reaction plate was centrifuged for 10 seconds to draw liquid to the bottom of the wells before being sealed with a Breath-Easy sealing membrane. The reaction plate was placed into a plate reader, heated to 45°C to accelerate eumelanin production and prevent potential cell growth from affecting optical density readings. To measure cell density in the reaction plate, an initial measurement at OD_600_ is taken, after which OD_405_ measurements are taken every 10 minutes for 12 hours whilst the plate is shaken at high speed.

### Wettability experiments

*K. rhaeticus* _*p*_*tyr1* was inoculated into a 24 well deep well plate, with each well containing 5 mL of HS-glucose media, with 0.5 g/L tyrosine, 10 μM CuSO_4_ and 340 μg/mL chloramphenicol. After 7 days at 30°C, pellicles were harvested. Eumelanin production was initiated, by placing the harvested pellicles into eumelanin development buffer. A set of pellicles were held back from eumelanin production and placed into an acetate buffer containing 0.5 g/L tyrosine, 10 μM CuSO_4_ at pH 3.6 to act as a negative control. Melanated and unmelanated pellicles were sterilised by placing them in 70% ethanol overnight. Pellicles were then washed in distilled water to remove left-over ethanol and salt. Pellicles were then dried flat using a heated press set to 120°C and 1 ton of pressure. To facilitate this drying and prevent the pellicles sticking to the press, pellicles were sandwiched between 3 layers of filter paper. Wettability tests were conducted using a KRUSS EasyDrop with 1 μL of water. Each contact angle measurement was derived from the average contact angle from 10 back-to-back water drop images taken within 10 seconds of drop contact with the pellicle surface.

### Tensile strength experiments

*K. rhaeticus* _*p*_*tyr1* was inoculated into 15cm square petri dishes containing 50 mL of HS-glucose media, with 0.5 g/L tyrosine, 10 μM CuSO_4_ and 340 μg/mL chloramphenicol. After 7 days at 30°C, pellicles were harvested and cut in half. One half was placed into an eumelanin development buffer to initiate eumelanin production and the other half into an acetate buffer containing 0.5 g/L tyrosine, 10 μM CuSO_4_ at pH 3.6 to prevent eumelanin production. After 24 hours shaking at 30°C, melanated and unmelanated pellicles were removed from their respective buffers and sterilised in a 70% ethanol solution overnight. Pellicles were then washed in distilled water to remove ethanol and salts left over from the eumelanin development processes. Pellicles were then dried flat using a heated press set to 120°C and 1 ton of pressure. Dog bone test specimens 35 mm long were cut out of the dried cellulose using a Zwick ZCP 020 manual cutting press. Pellicle specimen ends reinforced with card using Everbuild Stick 2 superglue. Dots were marked on the surface of each specimen for the optical measurement of displacement. A silver pen was used to dot melanated cellulose to generate the necessary contrast for optical measurement of displacement. Tensile tests were conducted with a Deben Microtest Tensile Stage using a load cell of 200 N and crosshead speed of 0.5 mm min-1.

### Scanning electron microscopy

The unmelanated pellicle was prepared by placing it into an acidic acetate buffer at pH 3.6, which prevented eumelanin synthesis and incubated in identical conditions to the melanated pellicle in the eumelanin development buffer bath. Melanated and unmelanated pellicles were prepared for SEM through the following steps. Unsterilised pellicles were placed in a 20% ethanol solution and shaken gently for 1 hour before being removed and placed into a 40% ethanol solution and shaken gently. This process was repeated for 60%, 80% and 100% ethanol solutions to ensure the maximum replacement of water with ethanol from the cellulose matrix. Pellicles were then flash frozen in liquid nitrogen and freeze dried until completely dry. The fully dried pellicles were then gold coated and imaged under high tension with a JSM6400 scanning electron microscope.

### Light microscopy

*K. rhaeticus* _*p*_*tyr1* was inoculated into 3 ml of HS-glucose media containing 2% (v/v) cellulase and 340 μg/mL chloramphenicol and grown shaking at 30°C until turbid. The turbid culture was then pelleted by centrifugation, washed with 1 mL PBS and split into two separate 1.5 centrifuge tubes. The cells were then pelleted again. One pellet was resuspended with 500 μL eumelanin development buffer to initiate eumelanin production and the other pellet resuspended with 500 μL PBS to keep the cells unmelantated. The cells were incubated over 24 hours at 30°C by which point the tube containing the cells in eumelanin development buffer had turned black. To prepare the microscope slides, 1 μl of melanated and unmelanated *K. rhaeticus* _*p*_*tyr1* cells were placed on separate 1% agar pads and imaged on a Nikon Ti-EX1 invert microscope. Cells were imaged in bright field with no phase contrast to accurately represent the shade of the cells.

### Patterning *mCherry* expression in a *K. rhaeticus* _p_Opto-T7RNAP*(563-F2)-*mCherry* pellicle

A custom projection rig was built to project light onto the growing pellicle (extended data: Fig. 4b). This held an acetate transparency that contained various components that would test the quality of the patterning in the pellicle. The image transparency was designed in Adobe Illustrator and printed on an HP LaserJet 500 MFP M570. Four acetate transparencies were stacked a top each other to form the final transparency. This was then sealed between glass slides and secured to the upper laboratory loop clamp. The pellicle container was sterilized and filled with 500 mL of HS-glucose media, containing 0.1 % (w/v) arabinose, 1% (v/v) ethanol and 170 μg/mL chloramphenicol. The media was then inoculated with a 1 mL *K. rhaeticus* _p_Opto-T7RNAP*(563-F2)-*mCherry* glycerol aliquot and a glass lid was placed on top of the container. This glass lid was warmed before placement to prevent condensation forming on it and distorting the projection. The LED lamp was then turned on, the lens shuttered with a piece of black card. After 3 days at ∼30°C, a thin pellicle had formed. The lens was uncovered and the image from the transparency focused on the pellicle. Once the pellicle had been exposed to the projected image for 3 days, it was harvested and scanned using a FLA-5000 Fluorescence scanner (Fujifilm). Image analysis was conducted using the OpenCV Python library.

### Patterning *tyr1* expression in a *K. rhaeticus* Opto-T7RNAP(563-F1)-*tyr1* pellicle

A custom rig using a commercial LED projector (ViewSonic M1) was built to project light onto the growing pellicle (Extended data: Fig. 4e). The rig was draped with blackout fabric to remove outside light. A timelapse image was designed in Adobe Illustrator to test how long a given pellicle would need to be exposed to light before an identifiable change in pigmentation could be observed. In this image, Blue is represented by an RGB value of 0,0,255, cyan by 0,255,255, white by 255,255,255 and black by 0,0,0. The pellicle container was sterilized and filled with 1 L of coconut water media, containing 1 % (w/v) arabinose, 0.5 g/L L-tyrosine, 10 μM CuSO_4_, 1% (v/v) ethanol and 200 μg/mL spectinomycin. The media was then inoculated with a 1 mL *K. rhaeticus* Opto-T7RNAP(563-F1)-t*yr1* glycerol aliquot and the culture container covered with foil. After 8 days at room temperature (∼20°C), a thin pellicle had formed. The foil was then removed, the projector focused on the surface of the pellicle, and the 80-hour video started. After 80 hours, the pellicle was harvested and placed into a 300×400 mm Eurobox containing 2 L of concentrated eumelanin development buffer and left to develop in stationary conditions at 30°C until a discernible pattern could be identified. The pellicle was then washed in dH_2_0 to remove eumelanin that had not accumulated within the pellicle. Densitometry scans of the pellicle were taken using an Amersham Typhoon scanner (GE), set to the Digi-blue digitalisation setting.

### Characterising *mCherry* expressing optogenetic strains

*K. rhaeticus* Opto-T7RNAP strains carrying the _*p*_*T7-mCherry* plasmid and *K. rhaeticus* _P_Opto-T7RNAP*(563-F2)-*mCherry* were cultured, in darkness, shaking in 3 mL of HS-glucose media with 2% cellulase, containing either spectinomycin at 200 μg/mL or chloramphenicol at 340 μg/ml depending on the plasmid. When all cultures had become turbid, the OD_600_ was measured and cultures were all either diluted or concentrated to an OD_600_ of 1, before being inoculated 1:10 into a 96-well deep well plate containing 270 μL HS-glucose media with 2% cellulase and either 0, 1, 10 or 100 mg/ml of arabinose. Where appropriate, spectinomycin at 200 μg/mL and chloramphenicol at 340 μg/ml were added to the wells. After 18 hours shaking growth at 30°C in darkness, cells were split across two clear 96 well plates, diluted 1:2 into fresh media with a matching arabinose concentration. One plate was placed onto a shaker under a blue LED flood light and the other plate wrapped in foil and placed on the same shaker. Both plates were sealed with Breath-Easy sealing membrane. After 6 hours in the two lighting conditions at 30°C and fast shaking, the cells were placed into a plate reader and red fluorescence in each well was measured using Ex:590 nm and Em:645 nm as well as cell density at OD_600_.

### Characterising *tyr1* expressing optogenetic strains

The Opto-T7RNAP *K. rhaeticus* strains carrying the _*p*_*T7-tyr1* plasmid and *K. rhaeticus* _P_Opto-T7RNAP(563-F1)-*tyr* were cultured in the same manner as the mCherry strains — with the exception that the HS-glucose was supplemented with 0.5 g/L tyrosine and 10 μM CuSO_4_. The approach to exposing the cells to blue light was also the same as the mCherry strains, except, after the 6 hours exposure time, the two plates were entered into the eumelanin production assay procedure. The two plates were placed onto the OT-2 deck and samples from both plates were mixed with eumelanin development buffer in a 384 well reaction plate. Each well in the two 96 well plates was sampled twice in the 384 reaction plate to give two technical replicates for each well. These two replicates were then averaged during analysis.

## Acknowledgements

We wish to thank Andreas Kamolz and Alice Potts at Modern Synthesis for construction and photography of the wallet prototype, Ed Tritton for photography of the shoe upper, Ben Reeve at Modern Synthesis for culture scale-up advice and manuscript feedback, Tanya Tschirhart at the U.S. Naval Research Laboratory for informative dialogue on the Tyr1 tyrosinase and Carole Collet at Central Saint Martins, University of the Arts London, for creating the cross-disciplinary environment that seeded the ideas for this work.

## References

1. Sadowski, M., Perkins, L. & McGarvey, E. Roadmap to Net Zero: Delivering Science-Based Targets in the Apparel Sector. World Resour. Inst. 1–40 (2021). doi:10.46830/wriwp.20.00004

2. Tkaczyk, A., Mitrowska, K. & Posyniak, A. Synthetic organic dyes as contaminants of the aquatic environment and their implications for ecosystems: A review. Sci. Total Environ. 717, 137222 (2020).

3. Liu, J. et al. Microfiber pollution: an ongoing major environmental issue related to the sustainable development of textile and clothing industry. Environ. Dev. Sustain. 2021 238 23, 11240–11256 (2021).d

4. Jones, M., Gandia, A., John, S. & Bismarck, A. Leather-like material biofabrication using fungi. Nat. Sustain. 2020 41 4, 9–16 (2020).

5. Meyer, M., Dietrich, S., Schulz, H. & Mondschein, A. Comparison of the Technical Performance of Leather, Artificial Leather, and Trendy Alternatives. Coatings 2021, Vol. 11, Page 226 11, 226 (2021).

6. Tang, T. C. et al. Materials design by synthetic biology. Nat. Rev. Mater. 2020 64 6, 332–350 (2020).

7. Gilbert, C. & Ellis, T. Biological Engineered Living Materials: Growing Functional Materials with Genetically Programmable Properties. ACS Synth. Biol. 8, 1–15 (2019).

8. Ryngajłło, M., Kubiak, K., Jędrzejczak-Krzepkowska, M., Jacek, P. & Bielecki, S. Comparative genomics of the Komagataeibacter strains—Efficient bionanocellulose producers. Microbiologyopen 8, (2019).

9. Huang, Y. et al. Recent advances in bacterial cellulose. Cellulose 21, 1–30 (2014).

10. Vazquez, A., Foresti, M. L., Cerrutti, P. & Galvagno, M. Bacterial Cellulose from Simple and Low Cost Production Media by Gluconacetobacter xylinus. J. Polym. Environ. 21, 545–554 (2013).

11. Jozala, A. F. et al. Bacterial cellulose production by Gluconacetobacter xylinus by employing alternative culture media. Appl. Microbiol. Biotechnol. 99, 1181–1190 (2015).

12. Chawla, P. R., Bajaj, I. B., Survase, S. A. & Singhal, R. S. Microbial cellulose: Fermentative production and applications. Food Technology and Biotechnology 47, 107–124 (2009).

13. Ul-Islam, M., Khan, T. & Park, J. K. Water holding and release properties of bacterial cellulose obtained by in situ and ex situ modification. Carbohydr. Polym. 88, 596–603 (2012).

14. Lee, K.-Y., Buldum, G., Mantalaris, A. & Bismarck, A. More Than Meets the Eye in Bacterial Cellulose: Biosynthesis, Bioprocessing, and Applications in Advanced Fiber Composites. Macromol. Biosci. 14, 10–32 (2014).

15. Czaja, W., Krystynowicz, A., Bielecki, S. & Brown, R. M. Microbial cellulose--the natural power to heal wounds. Biomaterials 27, 145–51 (2006).

16. Portela, R., Leal, C. R., Almeida, P. L. & Sobral, R. G. Bacterial cellulose: a versatile biopolymer for wound dressing applications. Microbial Biotechnology 12, 586–610 (2019).

17. Vandamme, E. J., De Baets, S., Vanbaelen, A., Joris, K. & De Wulf, P. Improved production of bacterial cellulose and its application potential. Polym. Degrad. Stab. 59, 93–99 (1998).

18. Gwon, H. et al. A safe and sustainable bacterial cellulose nanofiber separator for lithium rechargeable batteries. Proc. Natl. Acad. Sci. U. S. A. 116, 19288–19293 (2019).

19. Jiang, F., Yin, L., Yu, Q., Zhong, C. & Zhang, J. Bacterial cellulose nanofibrous membrane as thermal stable separator for lithium-ion batteries. J. Power Sources 279, 21–27 (2015).

20. Huang, C. et al. Composite nanofiber membranes of bacterial cellulose/halloysite nanotubes as lithium ion battery separators. Cellulose 26, 6669–6681 (2019).

21. García, C. & Prieto, M. A. Bacterial cellulose as a potential bioleather substitute for the footwear industry. Microb. Biotechnol. 12, 582 (2019).

22. Gilbert, C. et al. Living materials with programmable functionalities grown from engineered microbial co-cultures. Nat. Mater. (2021). doi:10.1038/s41563-020-00857-5

23. Florea, M. et al. Engineering control of bacterial cellulose production using a genetic toolkit and a new cellulose-producing strain. Proc. Natl. Acad. Sci. 113, E3431–E3440 (2016).

24. Teh, M. Y. et al. An Expanded Synthetic Biology Toolkit for Gene Expression Control in Acetobacteraceae. ACS Synth. Biol. 8, 708–723 (2019).

25. Goosens, V. J. et al. Komagataeibacter tool kit (KTK): a modular cloning system for multigene constructs and programmed protein secretion from cellulose producing bacteria. bioRxiv 2021.06.09.447691 (2021). doi:10.1101/2021.06.09.447691

26. Yadav, V. et al. Novel in vivo-degradable cellulose-chitin copolymer from metabolically engineered Gluconacetobacter xylinus. Appl. Environ. Microbiol. 76, 6257–65 (2010).

27. Takahama, R., Kato, H., Tajima, K., Tagawa, S. & Kondo, T. Biofabrication of a Hyaluronan/Bacterial Cellulose Composite Nanofibril by Secretion from Engineered Gluconacetobacter. Biomacromolecules 22, 4709–4719 (2021).

28. Walker, K. T., Goosens, V. J., Das, A., Graham, A. E. & Ellis, T. Engineered cell-to-cell signalling within growing bacterial cellulose pellicles. Microb. Biotechnol. 12, 611–619 (2019).

29. Yao, J., Dou, C., Wei, S. & Zheng, M. Using ecological reducing agents instead of sodium sulphide in dyeing with CI Sulphur Black 1. Color. Technol. 131, 379–383 (2015).

30. Božič, M. & Kokol, V. Ecological alternatives to the reduction and oxidation processes in dyeing with vat and sulphur dyes. Dye. Pigment. 76, 299–309 (2008).

31. Glass, K. et al. Direct chemical evidence for eumelanin pigment from the Jurassic period. Proc. Natl. Acad. Sci. U. S. A. 109, 10218–23 (2012).

32. Ito, S. A Chemist’s View of Melanogenesis. Pigment Cell Res. 16, 230–236 (2003).

33. Brenner, M. & Hearing, V. J. The protective role of melanin against UV damage in human skin. Photochemistry and Photobiology 84, 539–549 (2008).

34. Vahidzadeh, E., Kalra, A. P. & Shankar, K. Melanin-based electronics: From proton conductors to photovoltaics and beyond. Biosensors and Bioelectronics 122, 127–139 (2018).

35. Turick, C. E., Ekechukwu, A. A., Milliken, C. E., Casadevall, A. & Dadachova, E. Gamma radiation interacts with melanin to alter its oxidation-reduction potential and results in electric current production. Bioelectrochemistry 82, 69–73 (2011).

36. Mostert, A. B. et al. Role of semiconductivity and ion transport in the electrical conduction of melanin. Proc. Natl. Acad. Sci. U. S. A. 109, 8943–8947 (2012).

37. Pacelli, C. et al. Melanin is effective in protecting fast and slow growing fungi from various types of ionizing radiation. Environ. Microbiol. 19, 1612–1624 (2017).

38. Gustavsson, M., Hörnström, D., Lundh, S., Belotserkovsky, J. & Larsson, G. Biocatalysis on the surface of Escherichia coli: Melanin pigmentation of the cell exterior. Sci. Rep. 6, 1–9 (2016).

39. Wang, Z. et al. Melanin Produced by the Fast-Growing Marine Bacterium Vibrio natriegens through Heterologous Biosynthesis: Characterization and Application. Appl. Environ. Microbiol. 86, 1–18 (2019).

40. Chávez-Béjar, M. I. et al. Metabolic engineering of Escherichia coli to optimize melanin synthesis from glucose. Microb. Cell Fact. 12, 108 (2013).

41. Lin, W.-P. et al. Effect of melanin produced by a recombinant Escherichia coli on antibacterial activity of antibiotics. J. Microbiol. Immunol. Infect. 38, 320—326 (2005).

42. Shuster, V. & Fishman, A. Isolation, Cloning and Characterization of a Tyrosinase with Improved Activity in Organic Solvents from Bacillus megaterium. J. Mol. Microbiol. Biotechnol. 17, 188–200 (2009).

43. Nakano, S. & Ebisuya, H. Physiology of Acetobacter and Komagataeibacter spp.: Acetic Acid Resistance Mechanism in Acetic Acid Fermentation. in Acetic Acid Bacteria 223–234 (Springer Japan, 2016). doi:10.1007/978-4-431-55933-7_10

44. Masaoka, S., Ohe, T. & Sakota, N. Production of cellulose from glucose by Acetobacter xylinum. J. Ferment. Bioeng. 75, 18–22 (1993).

45. Hestrin, S. & Schramm, M. Synthesis of cellulose by Acetobacter xylinum. 2. Preparation of freeze-dried cells capable of polymerizing glucose to cellulose. Biochem. J. 58, 345–352 (1954).

46. Settembre, E. C., Chittuluru, J. R., Mill, C. P., Kappock, T. J. & Ealick, S. E. Acidophilic adaptations in the structure of Acetobacter aceti N 5-carboxyaminoimidazole ribonucleotide mutase (PurE). Acta Crystallogr. Sect. D Biol. Crystallogr. 60, 1753–1760 (2004).

47. Phisalaphong, M. & Chiaoprakobkij, N. Applications and Products—Nata de Coco. in Bacterial NanoCellulose. CRC Press 143–155 (2016).

48. Baumschlager, A., Aoki, S. K. & Khammash, M. Dynamic Blue Light-Inducible T7 RNA Polymerases (Opto-T7RNAPs) for Precise Spatiotemporal Gene Expression Control. ACS Synth. Biol. 6, 2157–2167 (2017).

49. Liu, Z. et al. Programming bacteria with light-sensors and applications in synthetic biology. Front. Microbiol. 9, 2692 (2018).

50. Teh, M. Y. et al. An Expanded Synthetic Biology Toolkit for Gene Expression Control in Acetobacteraceae. ACS Synth. Biol. 8, 708–723 (2019).

51. Ito, S. Optimization of Conditions for Preparing Synthetic Pheomelanin. Pigment Cell Res. 2, 53–56 (1989).

52. Lee, J. et al. Production of Tyrian purple indigoid dye from tryptophan in Escherichia coli. Nat. Chem. Biol. 2020 171 17, 104–112 (2020).

53. Fabara, A. N. & Fraaije, M. W. Production of indigo through the use of a dual-function substrate and a bifunctional fusion enzyme. Enzyme Microb. Technol. 142, 109692 (2020).

