## Extended Data for "Self-dyeing textiles grown from cellulose-producing bacteria with engineered tyrosinase expression"

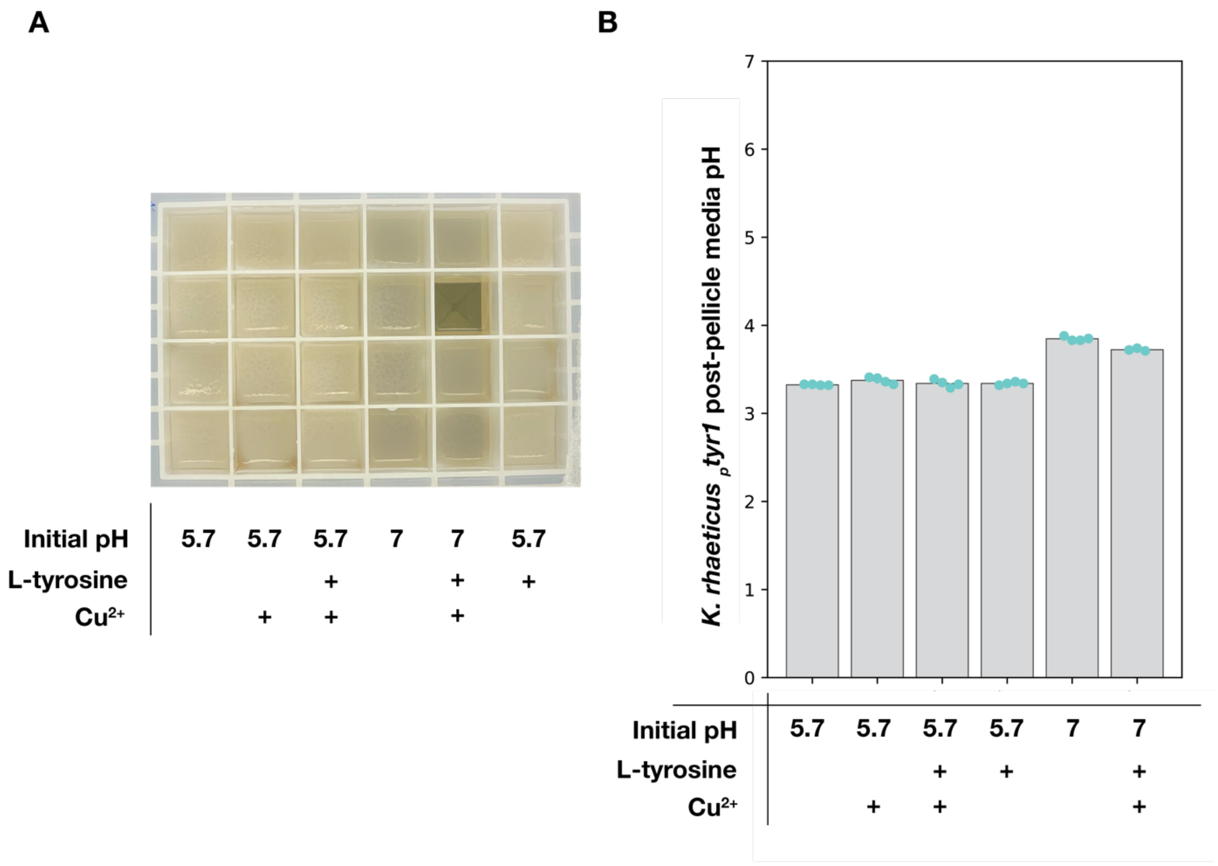


**Extended figure 1. Post-pellicle growth media pH of *K. rhaeticus* $_{P}$*tyr1*. A)** Pellicles produced after 7 days growth at 30°C in a 24 well deep well plate. Each plate column contains 4 replicated media compositions. Buffered HS-glucose media pH values before growth are noted beneath, as well as additional components required for melanin production. **B)** pH values of media underneath the pellicle after 7 days growth at 30°C. Data shown are average and standard deviation of 4 replicates, except for media set to pH 7 with L-tyrosine and Cu, where only 3 replicates produced pellicles.

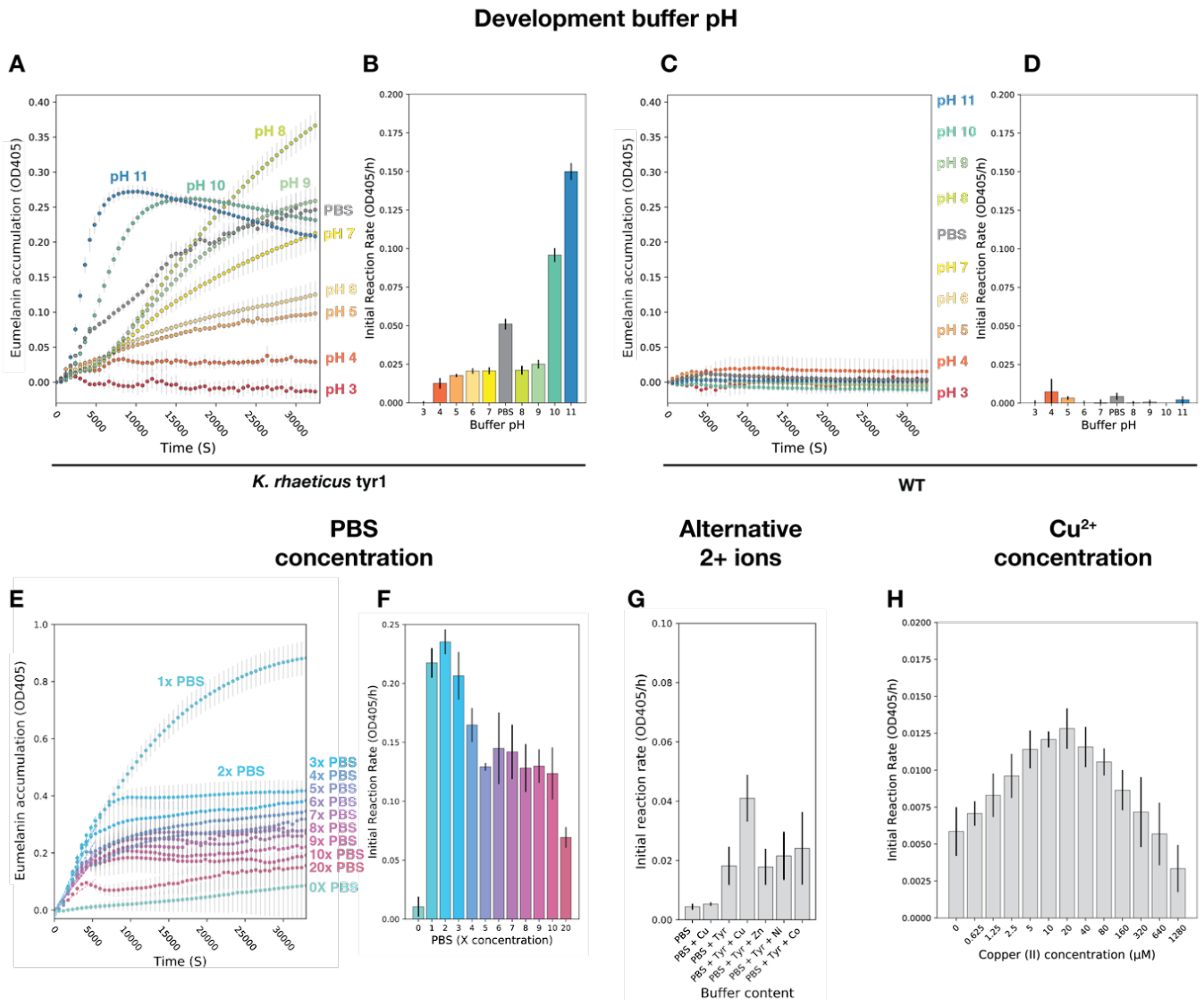


**Extended Figure 2. Characterisation of melanin production rates in differing melanin development buffer formulations. A)** A 10 hour time course experiment of eumelanin accumulation at $OD_{405}$ from *K. rhaeticus tyr1* in a range of pH values. *K. rhaeticus tyr1* cells were monitored in a citrate-phosphate-borate buffer containing 0.5 g/L L-tyrosine and 10 µM $CuSO_4$ set to various pH values between 3 and 11. Phosphate buffered saline (PBS) set to 7.4 containing 0.5 g/L L-tyrosine and 10 µM $CuSO_4$ is shown in grey. Error bars represent the standard deviation of 6 replicates. **B)** Average initial reaction rates of melanin accumulation from *K. rhaeticus tyr1* in buffers of differing pH values. Reaction rates are derived from the gradient of $OD_{405}$ over the first 140 minutes of the experiment. Colours used match those in panel A. Error bars represent the standard deviation of 6 replicates. **C - D)** 10 hour time course experiment of melanin accumulation and initial reaction rates of melanin accumulation from *K. rhaeticus* WT**.** Error bars represent standard deviation of 6 technical replicates. **E)** A 10 hour time course experiment of melanin accumulation at $OD_{405}$ from *K. rhaeticus tyr1* in differing concentrations of PBS. A PBS concentration of 0x corresponds to $ddH_20$ containing only melanin development substrates (L-tyrosine and $CuSO_4$) while a PBS concentration of 1x corresponds to 137 mM NaCl, 2.7 mM KCL, 10 mM $Na_2HPO_4$ and 1.8 mM $KH_2PO_4$. Melanin development substrates are maintained at 0.5 g/L tyrosine and 10 µM $CuSO_4$ for all PBS concentrations. Error bars represent standard deviation of 6 replicates. **F)** Average initial reaction rates of melanin accumulation from *K. rhaeticus tyr1* in differing PBS concentrations. Reaction rates are derived from the gradient of $OD_{405}$ over the first 140 minutes of the experiment**. G)** Average initial reaction rates of melanin accumulation from *K. rhaeticus tyr1* in buffers containing different metal (II) ions. Ions present in solution at 20µM. **H)** Average initial reaction rates of melanin accumulation from *K. rhaeticus tyr1* in buffers containing different Copper (II) concentration.

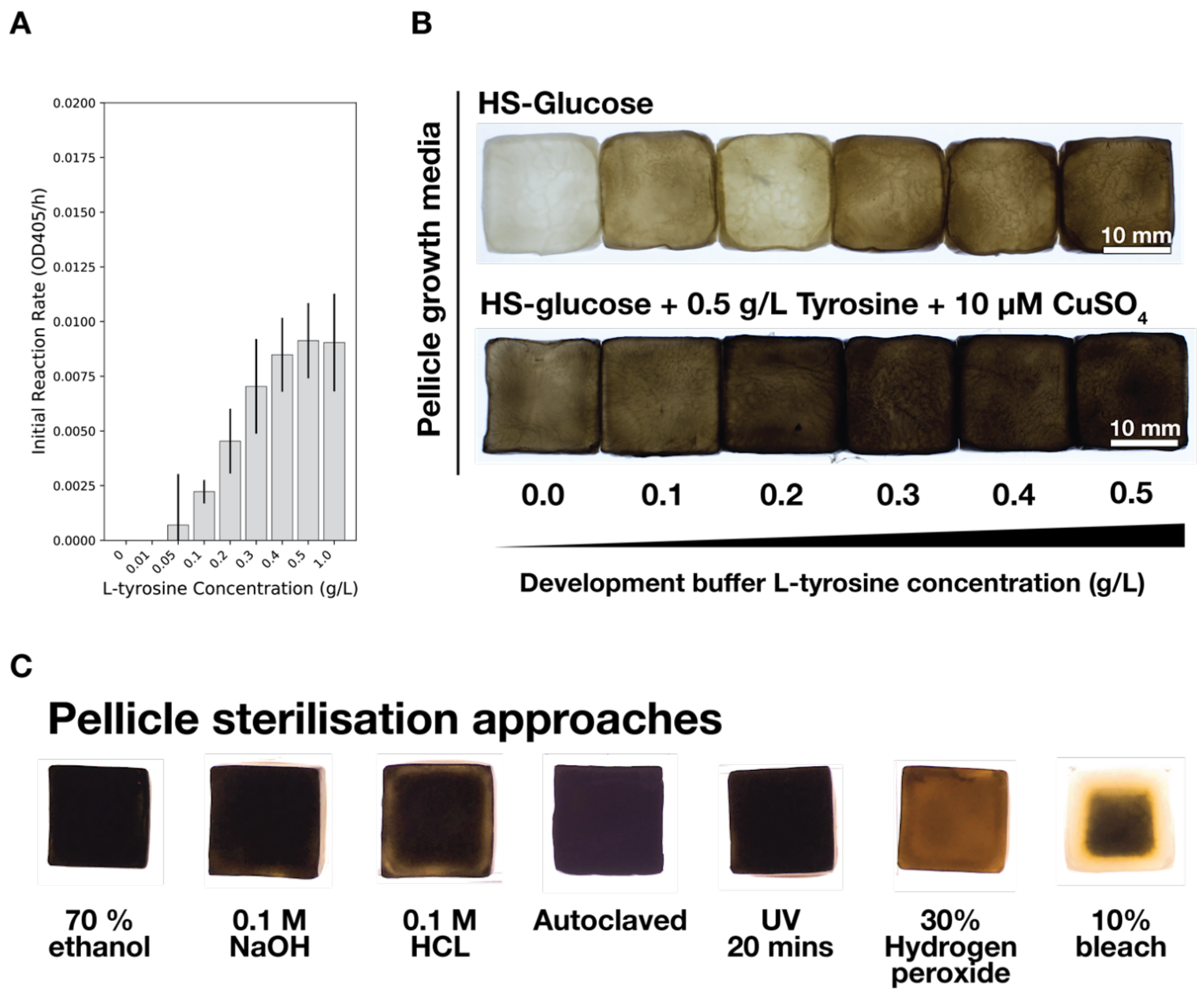


**Extended Figure 3. Melanin pigmentation strength and stability during sterilisation. A)** Average initial reaction rates of melanin accumulation from *K. rhaeticus tyr1* in PBS buffer with 10 µM $CuSO_4$ with differing concentrations of L-tyrosine. An L-tyrosine concentration of 1 g/L is above the maximum solubility of tyrosine in water at 25°C — $OD_{405}$ readings at this concentration and above become inaccurate due to suspended undissolved L-tyrosine. Error bars represent the standard deviation of 6 replicates and reaction rates are derived from the first 140 minutes of the reaction. **B)** *K. rhaeticus* $_{p}$*tyr1* pellicle pigmentation after 24 hours in a shaking melanin development bath at 30°C containing differing L-tyrosine concentration. The top row show pellicles from *K. rhaeticus* $_{p}$*tyr1* grown in HS-glucose media without L-tyrosine and $CuSO_4$ whilst the bottom row show pellicles grown in HS-glucose media with L-tyrosine and $CuSO_4$. **C)** Effects of various sterilisation methods on melanin stability in melanated pellicles from *K. rhaeticus* $_{p}$*tyr1*. Pellicles had been bathed in development buffer containing 0.5 g/L L-tyrosine and 10 µM $CuSO_4$ for 24 hours. Pellicles were exposed to sterilising conditions for 2 hours, except gel doc UV light, to which the pellicle was exposed to for 20 minutes.

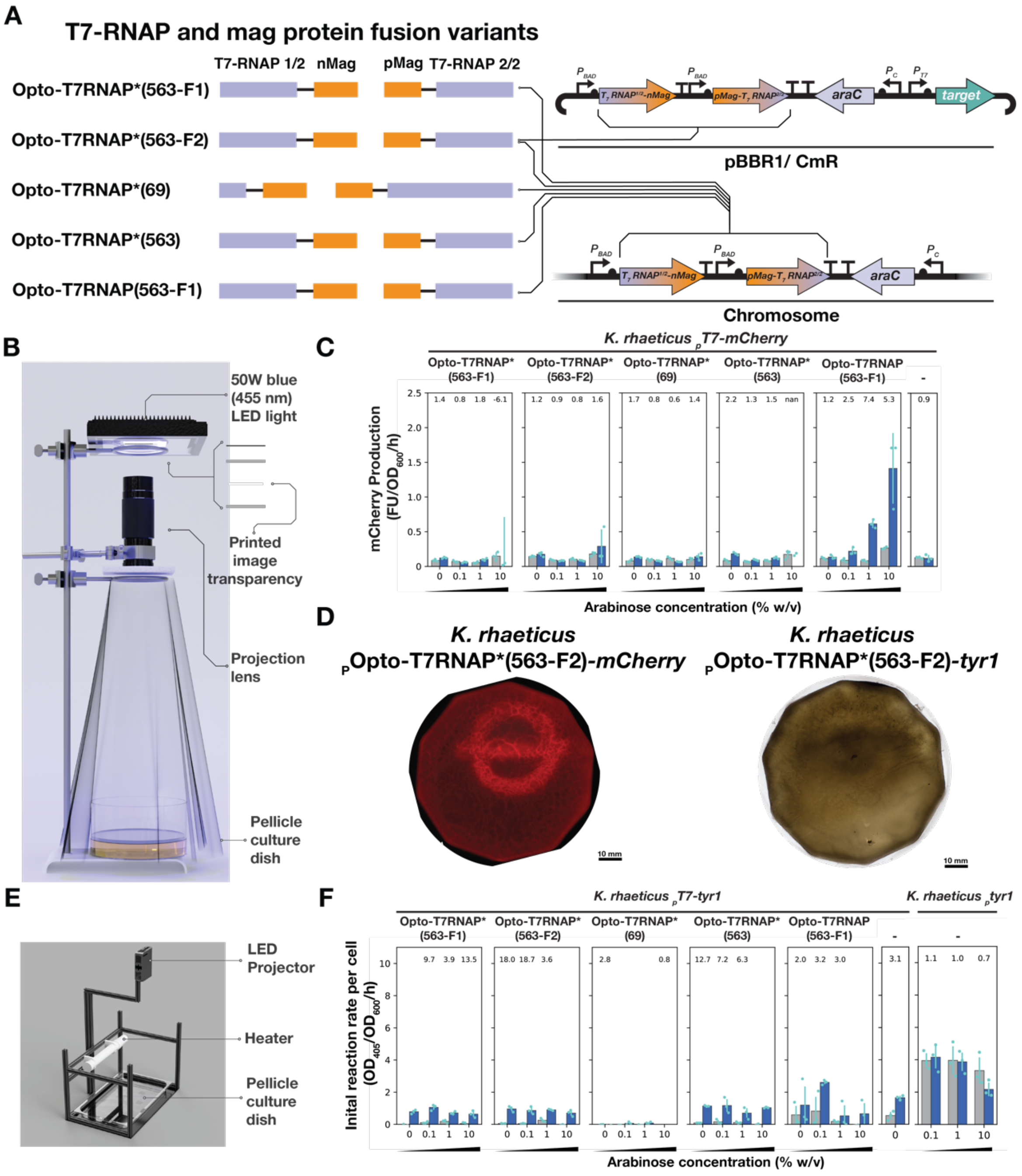


**Extended figure 4. Additional optogenetic data.** **A)** 5 variants of the opto-T7RNAPs system were integrated into the *K. rhaeticus* chromosome. Only Opto-T7RNAP*(563-F2) was used to create the plasmid-based arrangement. **B)** A rig for projecting light onto a growing pellicle. Blue-light from LED floodlight passes through an acetate transparency, on which is a printed image. The light that passes through this image is then focused and projected beneath onto the growing pellicle. Overall assembly is 620 mm tall. **C)** Blue-light response from all tested integrated constructs that were transformed with an *mCherry* target gene plasmid. Bars (blue = exposed, grey = unexposed) show average increase in red fluorescence after 6 hours normalised by $OD_{600}$. Error bars show standard deviation of 3 biological replicates placed on the same plate. Fold difference between exposed and unexposed cells is shown at top. **D)** Comparison of *K. rhaeticus* $_{p}$Opto-T7RNAP*(563-F2)-*mCherry* pellicle fluorescence scan and $_{p}$Opto-T7RNAP*(563-F2)-*tyr1* pellicle photograph. Both pellicles are produced using identical methods. **E)** Commercial projector-based assembly for patterning blue light on to a growing pellicle **F)** Characterisation data from tested genome-integrated constructs that were transformed with a *tyr1* target gene plasmid. Fold difference is given above the bars, but not shown when rates are 0 or below.
