## Supplementary Materials for "Self-dyeing textiles grown from cellulose-producing bacteria with engineered tyrosinase expression"

### S1. Strains used in this study

| **Strains** | **Description** | **Reference** |
|---|---|---|
| *K. rhaeticus* iGEM | Strain of bacterial cellulose producing bacteria isolated from a kombucha tea SCOBY | Florea *et al.* |
| *K. rhaeticus* $_{p}$*tyr1* | Constitutive production of Tyr1 protein. Possess chloramphenicol resistance | This study |
| *K. rhaeticus tyr1* | Constitutive production of Tyr1 protein from integrated tyr1 gene at the *arsH* locus. Possess chloramphenicol resistance. | This study |
| *K. rhaeticus* $_{p}$Opto-T7RNAP*(563-F2)-mCherry | Constitutive AraC production confers arabinose sensitivity. Expression of both halves of the Opto-T7RNAP*(563-F1) protein are under pBad promoter control, and are upregulated in an increase of arabinose concentration. Blue light sensitivity in presence of arabinose leads to expression of mCherry under the T7 promoter. Possess chloramphenicol resistance. | This study |
| *K. rhaeticus* Opto-T7RNAP*(563-F1)-*mCherry* | Constitutive AraC production confers arabinose sensitivity. Expression of both halves of the Opto-T7RNAP*(563-F1) protein are under pBad promoter control, and are upregulated in an increase of arabinose concentration. Blue light sensitivity in presence of arabinose leads to expression of mCherry under the T7 promoter. Possess chloramphenicol and spectinomycin resistance. | This study |
| *K. rhaeticus* Opto-T7RNAP*(563-F2)-*mCherry* | Constitutive AraC production confers arabinose sensitivity. Expression of both halves of the Opto-T7RNAP*(563-F2) protein are under pBad promoter control, and are upregulated in an increase of arabinose concentration. Blue light sensitivity in presence of arabinose leads to expression of mCherry under the T7 promoter. Possess chloramphenicol and spectinomycin resistance. | This study |
| *K. rhaeticus* Opto-T7RNAP*(69)-*mCherry* | Constitutive AraC production confers arabinose sensitivity. Expression of both halves of the Opto-T7RNAP*(69) protein are under pBad promoter control, and are upregulated in an increase of arabinose concentration. Blue light sensitivity in presence of arabinose leads to expression of mCherry under the T7 promoter. Possess chloramphenicol and spectinomycin resistance. | This study |
| *K. rhaeticus* Opto-T7RNAP*(563)-*mCherry* | Constitutive AraC production confers arabinose sensitivity. Both halves of the Opto-T7RNAP*(563) gene are under pBad promoter control, and are upregulated in an increase of arabinose concentration. Blue light sensitivity in presence of arabinose leads to expression of mCherry under the T7 promoter. Possess chloramphenicol and spectinomycin resistance. | This study |
| *K. rhaeticus* Opto-T7RNAP(563-F1)-*mCherry* | Constitutive AraC production confers arabinose sensitivity. Both halves of the Opto-T7RNAP(563-F1) gene are under pBad promoter control, and are upregulated in an increase of | This study |

| | | |
|---|---|---|
| | arabinose concentration. Blue light sensitivity in presence of arabinose leads to expression of mCherry under the T7 promoter. Possess chloramphenicol and spectinomycin resistance. | |
| *K. rhaeticus* $_{p}$*T7-mCherry* | *mCherry* gene under the T7 promoter control. Does not contain a T7 polymerase gene. Possess spectinomycin resistance. | This study |
| *K. rhaeticus* $_{p}$Opto-T7RNAP*(563-F2)-*tyr1* | Constitutive AraC production confers arabinose sensitivity. Expression of both halves of the Opto-T7RNAP*(563-F1) protein are under pBad promoter control, and are upregulated in an increase of arabinose concentration. Blue light sensitivity in presence of arabinose leads to expression of Tyr1 under the T7 promoter. Possess chloramphenicol resistance. | This study |
| *K. rhaeticus* Opto-T7RNAP*(563-F1)-*tyr1* | Constitutive AraC production confers arabinose sensitivity. Expression of both halves of the Opto-T7RNAP*(563-F1) protein are under pBad promoter control, and are upregulated in an increase of arabinose concentration. Blue light sensitivity in presence of arabinose leads to expression of Tyr1 under the T7 promoter. Possess chloramphenicol and spectinomycin resistance. | This study |
| *K. rhaeticus* Opto-T7RNAP*(563-F2)-*tyr1* | Constitutive AraC production confers arabinose sensitivity. Expression of both halves of the Opto-T7RNAP*(563-F2) protein are under pBad promoter control, and are upregulated in an increase of arabinose concentration. Blue light sensitivity in presence of arabinose leads to expression of Tyr1 under the T7 promoter. Possess chloramphenicol and spectinomycin resistance. | This study |
| *K. rhaeticus* Opto-T7RNAP*(69)-*tyr1* | Constitutive AraC production confers arabinose sensitivity. Expression of both halves of the Opto-T7RNAP*(69) protein are under pBad promoter control, and are upregulated in an increase of arabinose concentration. Blue light sensitivity in presence of arabinose leads to expression of Tyr1 under the T7 promoter. Possess chloramphenicol and spectinomycin resistance. | This study |
| *K. rhaeticus* Opto-T7RNAP*(563)-*tyr1* | Constitutive AraC production confers arabinose sensitivity. Both halves of the Opto-T7RNAP*(563) gene are under pBad promoter control, and are upregulated in an increase of arabinose concentration. Blue light sensitivity in presence of arabinose leads to expression of Tyr1 under the T7 promoter. Possess chloramphenicol and spectinomycin resistance. | This study |
| *K. rhaeticus* Opto-T7RNAP(563-F1)-*tyr1* | Constitutive AraC production confers arabinose sensitivity. Both halves of the Opto-T7RNAP(563-F1) gene are under pBad promoter control, and are upregulated in an increase of arabinose concentration. Blue light sensitivity in presence of arabinose leads to expression of Tyr1 under the T7 promoter. Possess chloramphenicol and spectinomycin resistance. | This study |
| *K. rhaeticus* $_{p}$*T7-tyr1* | *tyr1* gene under the T7 promoter control. Does not contain a T7 polymerase gene. Possess spectinomycin resistance. | This study |

**S2. Plasmids used in this study**

| **Plasmid name** | **Description and construction** | **Reference** |
|---|---|---|
| $_p$Tyr1 | Plasmid constructed with KTK. pBBR1 origin of replication and chloramphenicol resistance cassette. J23104-B0034-Tyr1-L321P00 | This study |
| $_p$Tyr1_IV | Plasmid constructed with KTK. pUC19 origin of replication and ampicillin resistance cassette. J23104-B0034-Tyr1-L321P00 | This study |
| $_p$T7-mCherry | Plasmid constructed with Gibson cloning. pBBR1 origin of replication and spectinomycin resistance cassette. The *mCherry* coding sequence, RBS, terminator, and promoter were taken from pAB50 from Baumschlager *et al.* | This study |
| $_p$T7-tyr1 | Plasmid constructed with Gibson cloning. pBBR1 origin of replication and spectinomycin resistance cassette. Derived from $_p$T7-mCherry, using the same RBS, terminator, promoter but switching the mCherry CDS for the *tyr1* CDS. | This study |
| $_p$Opto-T7RNAP*(563-F2)-*mCherry* | Plasmid constructed with Gibson cloning. pBBR1 origin of replication and chloramphenicol resistance cassette. Both Opto-T7RNAP*(563-F2) genes were taken from pAB152 from baumschlager *et al.* | This study |
| $_p$Opto-T7RNAP*(563-F2)-*tyr1* | Plasmid constructed with Gibson cloning. pBBR1 origin of replication and chloramphenicol resistance cassette. Both Opto-T7RNAP*(563-F2) genes were taken from pAB152 from baumschlager *et al.* | This study |
| $_p$Opto-T7RNAP*(563-F1)_IV | Plasmid constructed with Gibson cloning. pUC19 origin of replication and Ampicillin resistance cassette. Both Opto-T7RNAP*(563-F1) genes were taken from pAB151 from baumschlager *et al.* | This study |
| $_p$Opto-T7RNAP*(563-F2)_IV | Plasmid constructed with Gibson cloning. pUC19 origin of replication and Ampicillin resistance cassette. Both Opto-T7RNAP*(563-F2) genes were taken from pAB152 from baumschlager *et al.* | This study |
| $_p$Opto-T7RNAP*(69)_IV | Plasmid constructed with Gibson cloning. pUC19 origin of replication and Ampicillin resistance cassette. Both Opto-T7RNAP*(69) genes were taken from pAB144 from baumschlager *et al.* | This study |
| $_p$Opto-T7RNAP*(563)_IV | Plasmid constructed with Gibson cloning. pUC19 origin of replication and Ampicillin resistance | This study |

| | | |
|---|---|---|
| | cassette. Both Opto-T7RNAP*(563) genes were taken from pAB150 from baumschlager *et al.* | |
| $_P$Opto-T7RNAP(563-F1)_IV | Plasmid constructed with Gibson cloning. pUC19 origin of replication and Ampicillin resistance cassette. Both Opto-T7RNAP(563-F1) genes were taken from pAB203 from baumschlager *et al.* | This study |

**S3. Tyr1 AA sequence.**

MGNKYRVRKNVLHLTDTEKRDFVRTVLILKEKGIYDRYIAWHGAAGKFHTPPGSDRNAAHMSSAFLPW
HREYLLRFERDLQSINPEVTLPYWEWETDAQMQDPSQSQIWSADFMGGNGNPIKDFIVDTGPFAAGRW
TTIDEQGNPSGGLKRNFGATKEAPTLPTRDDVLNALKITQYDTPPWDMTSQNSFRNQLEGFINGPQLH
NRVHRWVGGQMGVVPTAPNDPVFFLHHANVDRIWAVWQIIHRNQNYQPMKNGPFGQNFRDPMYPWNTT
PEDVMNHRKLGYVYDIELRKSKRSS*

**S4. *tyr1* DNA sequence.**

ATGGGCAATAAATACCGCGTGCGTAAGAATGTTCTGCACCTGACAGATACCGAGAAGCGTGACTTCGT
GCGCACTGTACTGATTTTGAAAGAGAAGGGCATTTACGATCGTTACATCGCATGGCACGGCGCCGCGG
GTAAGTTTCACACCCCGCCCGGTAGTGACCGTAACGCGGCGCACATGTCGAGTGCGTTCTTGCCTTGG
CACCGCGAATATCTGCTGCGCTTTGAGCGCGATCTGCAATCGATTAACCCTGAGGTGACTCTGCCGTA
CTGGGAGTGGGAAACCGATGCTCAAATGCAAGACCCTAGCCAGTCGCAGATCTGGAGCGCCGACTTCA
TGGGCGGCAATGGCAACCCAATTAAGGACTTCATTGTAGACACGGGCCCGTTCGCTGCCGGCCGTTGG
ACAACCATTGACGAGCAGGGTAACCCGTCAGGCGGCTTAAAGCGCAACTTCGGTGCGACTAAGGAAGC
CCCCACCCTGCCGACGCGCGACGACGTGCTGAACGCACTTAAGATTACCCAATACGACACCCCACCCT
GGGACATGACGTCCCAGAATAGTTTCCGCAACCAACTCGAGGGTTTCATCAATGGCCCGCAACTGCAT
AACCGTGTGCATCGCTGGGTCGGTGGCCAAATGGGTGTCGTCCCTACCGCGCCCAACGACCCGGTGTT
CTTCCTGCATCATGCGAACGTTGACCGCATCTGGGCCGTGTGGCAGATCATCCACCGCAACCAGAATT
ACCAACCAATGAAGAATGGCCCGTTCGGCCAGAATTTCCGTGACCCAATGTATCCATGGAACACCACG
CCTGAGGATGTAATGAATCACCGTAAACTGGGCTATGTTTATGACATCGAGTTGCGTAAGTCGAAGCG
CAGCTCTTGA

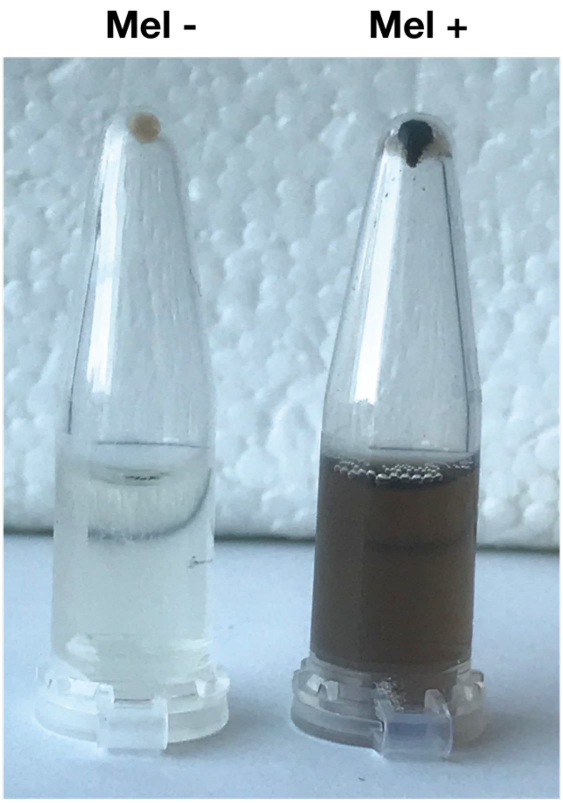


**S5. Pelleted unmelanated and melanated *K. rhaeticus* $_{p}$*tyr1* cells.** *K. rhaeticus* $_{p}$*tyr1* cells were grown in HS-glucose with 340 µg/ml chloramphenicol and 2 % (v/v) cellulase. Once turbid, cells were added to HS-glucose with 0.5 g/L L-tyrosine and 10 µM $CuSO_4$ with a citrate-phosphate buffer set to either pH 5.8 (Mel -) or pH 7 (Mel +). After 24 hours of shaking incubation at 30°C, 1 mL of cells from each culture were pelleted with centrifugation. Eumelanin pigmentation can be seen in both pellet and supernatant for cells that were exposed to HS-glucose media set to pH 7.

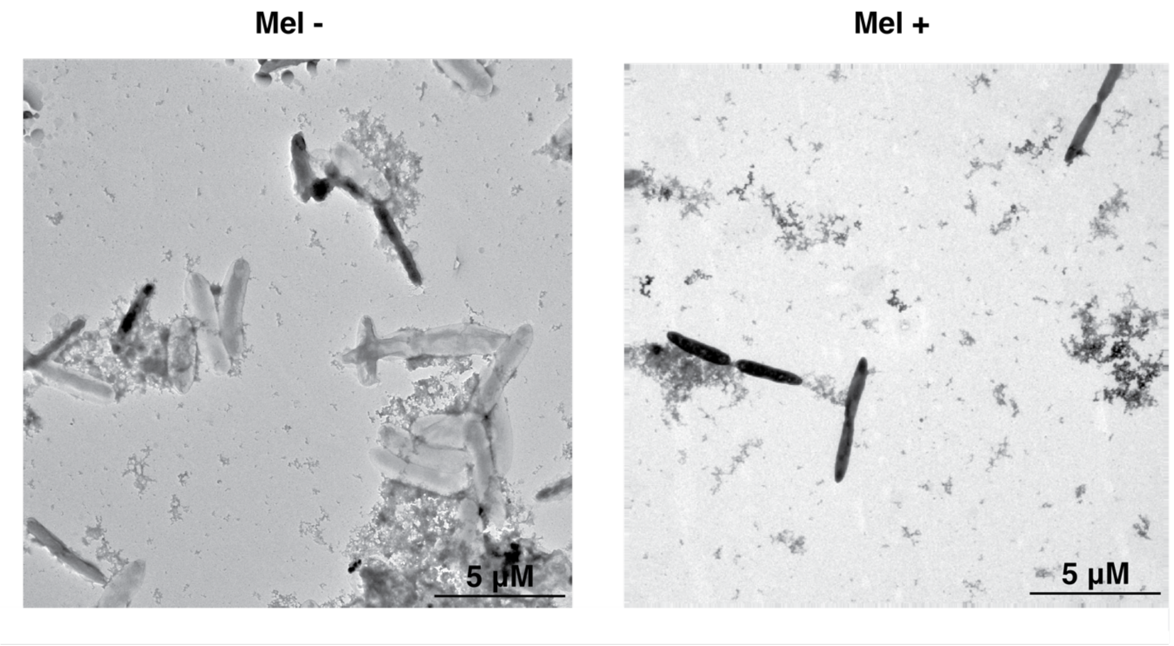


**S6. TEM microscopy of melanated and unmelanated *K. rhaeticus* $_{p}$*tyr1*.** Cells grown in HS-glucose media were washed in PBS, split into two separate tubes. PBS was replaced with either eumelanin development buffer or acetate buffer at pH 3.6 to produce melanated (mel +) and unmelanated cells (mel -) respectively. 2 µl samples were spotted onto 471 freshly glow discharged formvar/Carbon on 300 Mesh Nickel grids (Agar Scientific) and visualised with a FEI Tecnai G2 Spirit TWIN. Whilst cells were grown in 2% cellulase, globular matter seen in both images is likely to be incompletely digested cellulose.

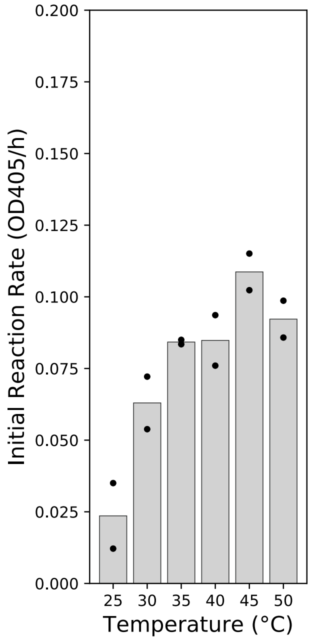


**S7. Initial reaction rate of eumelanin production from *K. rhaeticus tyr1* at a range of temperatures.** *K. rhaeticus tyr1* cells were grown in HS-glucose with 0.5 g/L L-tyrosine and 10 µM $CuSO_4$ before being washed and mixed with eumelanin development buffer. Cells were then distributed across a 96 well PCR plate using 50 uL per well. The plate was then placed into a heated block with a distributed range of temperatures. Every 20 minutes, a row of sample was removed, and placed on ice. After 120 minutes had passed eumelanin accumulation was measured at $OD_{405}$ for all temperatures and timepoints. Initial rate of reaction was calculated from the rate of eumelanin accumulation over 120 minutes. Two replicates were used for each temperature and bars show the average of these replicates.

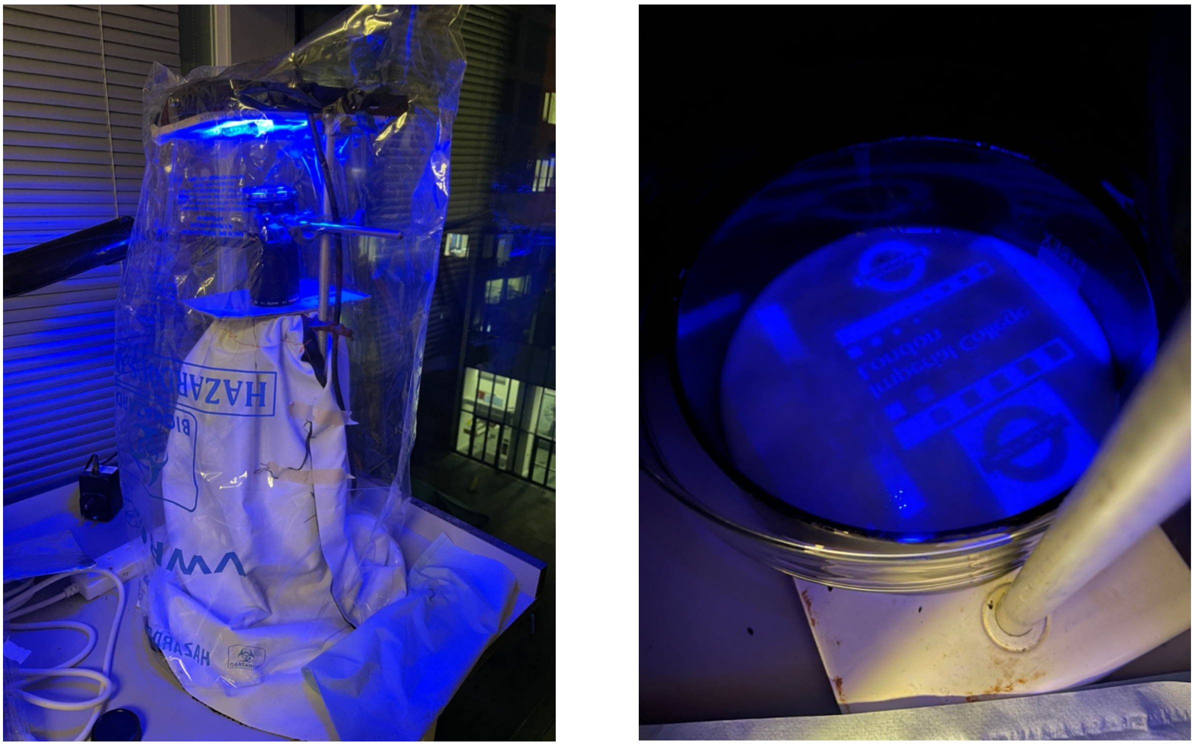

**S8. Photographs of optogenetic rig used to produce patterned pellicle from *K. rhaeticus* $_{p}$Opto-T7RNAP*(563-F2)-*mCherry*.** Image on the right shows the full assembly used to produce the patterned pellicle. Image on the right, shows the image transparency being projected on to the pellicle.

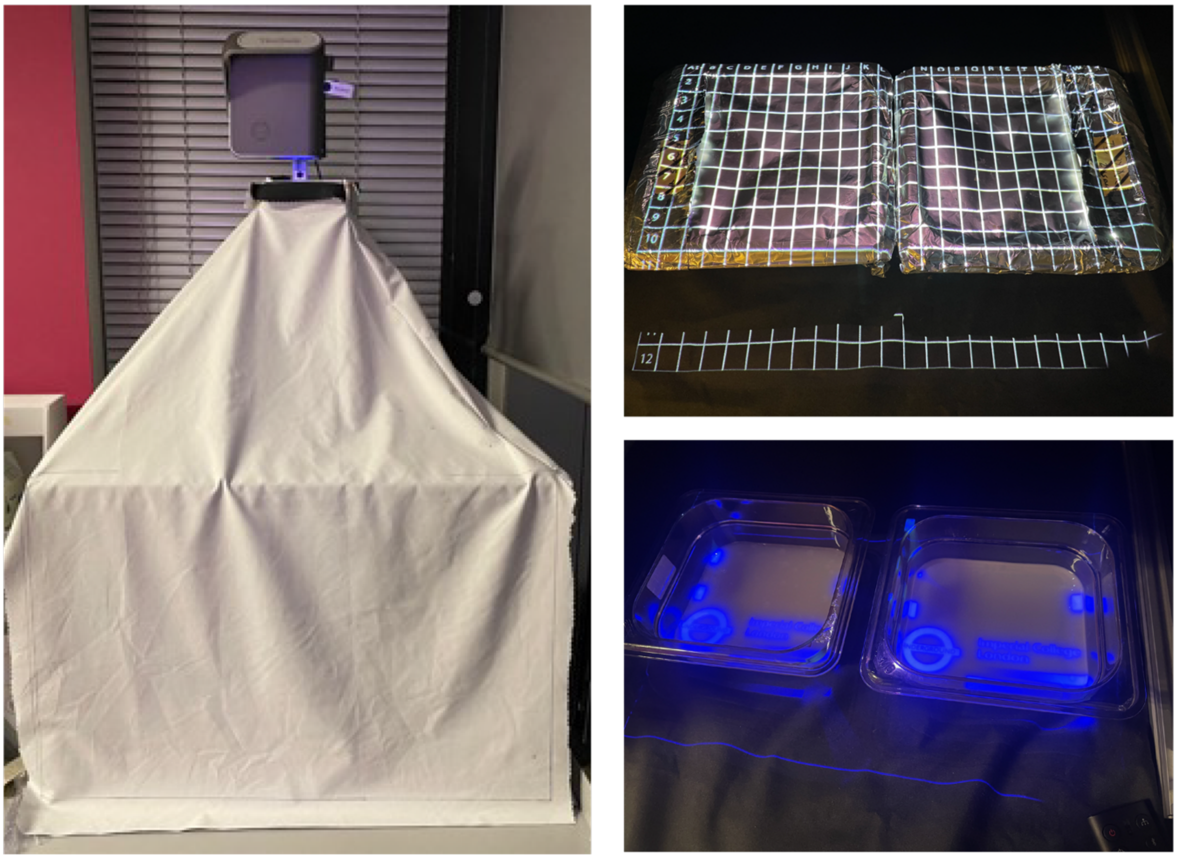

**S9. Photographs of optogenetic rig used to produce patterned pellicle from *K. rhaeticus*** $_{p}$**Opto-T7RNAP(563-F1)-*tyr1*.** Image on the right shows the full assembly used to produce the patterned *tyr1* pellicle. Assembly is coved in black out fabric to exclude light. Image on the top right, shows a grid being projected onto culture containers to aid in placement of image projections. Image on bottom right, shows to example images being projected onto a pair of pellicles.
